# Weed Instance Segmentation from UAV ortho-mosaic Images based on Deep Learning

**DOI:** 10.1101/2024.08.13.607729

**Authors:** Chenghao Lu, Kang Yu

## Abstract

Weeds significantly impact agricultural production, and traditional weed control methods often harm soil health and the environment. This study aims to develop deep learning-based segmentation models in identifying weeds in potato fields captured by Unmanned Aerial Vehicle (UAV) orthophotos and to explore the effects of weeds on potato yield. UAVs were used to collect RGB data from potato fields, flying at an altitude of 10m, with Real-ESRGAN Super-Resolution (SR) enhancing image resolution. We applied the Segment Anything Model (SAM) to do semi-automatic annotation, followed by training the YOLOv8 and MASK-RCNN models for segmentation. Also we used ANOVA and linear regression to analyze the effects of weeds and nitrogen fertilizer on yield. Results showed that the detection accuracy mAP50 scores for YOLOv8 and Mask R-CNN were 0.902 and 0.920, respectively, with the Real-ESRGAN-enhanced model achieving 0.909. When multiple weed types were present, accuracy decreased to 0.86. The linear regression model, incorporating plant and weed coverage areas, explained 41.2% of yield variation (R^2^ = 0.412, p-value = 0.01). Both YOLOv8 and Mask R-CNN achieved high accuracy, with YOLOv8 converging faster. Real-ESRGAN reconstruction slightly improved accuracy. Different nitrogen fertilizer treatments had no significant effect on yield, while weed control treatments significantly impacted yield, showing the importance of precise weed mapping.. This study provides insights into weed segmentation and contributes to environmentally friendly precision weed control.

**Graphic Abstract:** 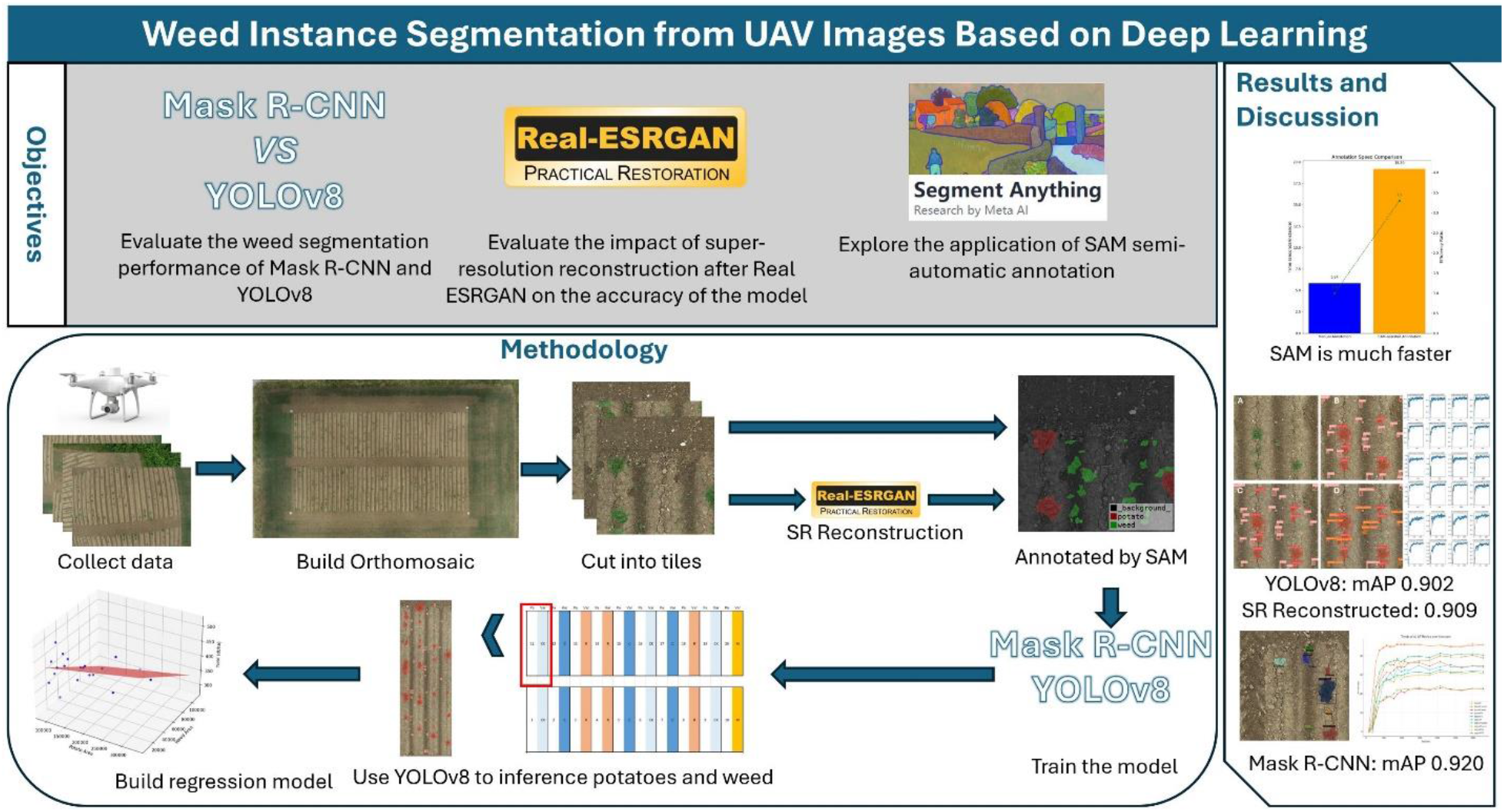

## 1 Introduction

Weeds proliferate in farmlands, gardens, and other human-managed environments, competing with crops for light, water, nutrients, and space. Due to their rapid growth and reproduction, weeds are challenging to eradicate and necessitate continuous management and control measures. Certain weed species threaten local ecosystems, invade and degrade natural environments, and negatively impact local plant populations and biodiversity (Kumar Rai and Singh, 2020). To manage weeds, farmers must invest additional labor, time, and resources in herbicides and mechanical weeding equipment. This increases crop production costs and may introduce harmful chemicals to the environment, causing long-term ecological impacts.

Precision weeding effectively reduces competition between crops and weeds, ensuring crops receive sufficient water, light, and nutrients (Riemens et al., 2022). This promotes healthy crop growth, increases yields, and improves crop quality while reducing labor and energy consumption, thereby lowering production costs. Minimizing chemical herbicide use helps protect farmland and surrounding environments, reduces pollution of water, soil, and air, and maintains field biodiversity and ecological balance, promoting the health and sustainability of agricultural ecosystems (Storkey and Westbury, 2007). Precision weeding employs various strategies and technologies to achieve effective weed management with minimal environmental impact and production costs (Shaner and Beckie, 2014). Advances in technology, such as artificial intelligence, machine learning, and machine vision, will enable more precise, efficient, and environmentally friendly weed control in the future.

Image segmentation divides an image into parts for easier analysis, using methods like thresholding, K-means clustering, edge detection, region growing, and deep learning with CNNs (Goyal and Punam, 2022). These techniques aid in object recognition, classification, and tracking. Image classification has advanced from simple color- and texture-based methods to deep learning models. Machine learning introduced feature-based classifiers like support vector machines (Chapelle et al., 1999). Recently, deep learning, especially CNNs, has transformed image classification, achieving high accuracy by learning features from data (Albelwi and Mahmood, 2016).

For instance, deep learning models have been increased utilized for weed detection, a major breakthrough in deep learning for weed detection has been the development of sophisticated architectures such as U-Net, Mask R-CNN, and YOLO (Guo et al., 2023). U-Net is a fully convolutional network (FCN) that is particularly effective for pixel-level classification tasks and is well suited for segmenting weeds from background and crops (Cui et al., 2024). Mask R-CNN extends the Faster R-CNN model by adding a branch that predicts segmentation masks, providing accurate localization and classification of weeds (He et al., 2017). YOLO is known for its real-time object detection capabilities, which balances speed and accuracy, making it ideal for applications that require immediate decision making (Redmon et al., 2016). For example, YOLOv5 is used to quickly detect maize seedlings in UAV images (Lu et al., 2024), the YOLOv5 model is deployed on NVIDIA Jetson nano and NVIDIA Jetson Orin devices to achieve real-time detection of crops (Nnadozie et al., 2024), and the YOLOWeeds ground image weed detection model developed based on YOLO (Dang et al., 2023). These applications have demonstrated the feasibility of YOLO for drone weed detection.

In the instance segmentation task of apple orchards, YOLOv8 demonstrated superior performance to Mask R-CNN, specifically, YOLOv8 achieved 92.9% precision and 97% recall in a single-class segmentation task, while Mask R-CNN achieved 84.7% precision and 88% recall (Sapkota et al., 2024). However, there has been no research comparing the capabilities of Mask R-CNN and YOLOv8 in segmenting weeds in UAV images. Li et al. trained a model based on a public weed dataset using eight popular segmentation algorithms, including Faster R-CNN, YOLOv3, YOLOv4, SSD, CenterNet, RetinaNet, EfficientDet, and YOLOX, achieving a maximum mAP of 79.63% (Li et al., 2022). Shahi et al. used UNet to train a segmentation model on the CoFly-WeedDB dataset based on UAV images of 5-meter flight height, with an accuracy of 88.20% (Shahi et al., 2023). Genze et al. released a manually annotated and expert-curated dataset of drone images, although the dataset used expensive cameras and had a GSD of 1mm, the UAV images were not corrected and were affected by motion blur, and their results showed that a UNet-like architecture with a ResNet-34 feature extractor achieved an F1 score of over 89% on the test set (Genze et al., 2022). However, most of these studies used UNet, but UNet requires high-resolution images, and the performance is limited in complex field environments and low-resolution images. The limitation of low resolution of UAV images is often only solved by reducing the flight altitude or by using more expensive cameras some studies.

Real-World Enhanced Super-Resolution Generative Adversarial Network (Real-ESRGAN) is an advanced image super-resolution technology based on deep learning, designed to enhance low-resolution images by improving their details to high-resolution quality (Wang et al., 2021). As an extension of ESRGAN, Real-ESRGAN is specifically optimized for real-world super-resolution tasks, capable of handling low-quality inputs, including compression artifacts, blur, and noise, to produce visually satisfying high-resolution images (Zhu et al., 2023). Their image enhancing techniques bring new possibilities of detecting small weeds from UAV image mosaic cover large fields. Yet, the feasibility of using the enhanced images in weed detection is still under explored. In addition, the enhance images might enable more precise and automated image annotations, and thus the deep leaning based image superesolution methods should be adapted to weed annotation and weed detection applications.

Here, our research question was can the image super resolution techniques improve the training data annotation and further improve the accuracy of deep learning models for weed segmentation? We hypothesize that Real-ESRGAN can help us improve the resolution of UAVs and thereby enable the SAM to realize semi-automatic annotation. The main objectives of the study are to: 1) Compare the weed segmentation performance of Mask R-CNN and YOLOv8, compare the performance of the YOLOv8 model in the presence of multiple types of weeds; 2) Evaluate the effects of super-resolution reconstruction after Real ESRGAN on the model performance of SAM semi-automatic annotation, as well as Mask RCNN and yolo on weed detection; and 3) verify the feasibility of applying the model for evaluating weed effects on potato yield.

## 2 Methodology

### 2.1 Experiment Site

The experimental site is located at the Agricultural Management Experiment Station of the Technical University of Munich in Dürnast, Freising, Germany (48^°^24’N, 11^°^42’E, altitude 485 m). The area of GHL is 570 m^2^, which we divided into 20 small areas of 3*m* ×6*m*. Each plot is planted with 4 rows of potatoes, the row spacing is 0.75m, the seeding density is 0.384 kg/m^2^, and the potato variety is Simonetta. On May 31, 2023, we carried out potato sowing in the GHL field, carried out fertilization treatment on June 6, carried out corresponding chemical weeding treatment on June 14, and carried out mechanical weeding treatment on July 12. The nitrogen fertilizer treatments include 75kg/ha and 150kg/ha. Weed treatments included controlled, chemical, robotic, and mechanical, with three replicates for each weed treatment.

### 2.2 UAV Images Collection

For UAV image collection, Ground Control Points (GCPs) are deployed at the four corners of the farmland. A ppm10xx GNSS Sensor (ppm GmbH, Penzberg, Germany) was used to obtain position and used for georeferencing. The drone used in this experiment is DJI Phantom 4 RTK (DJI, Shenzhen, China), The flight height was 10 m above ground level, and the flight speed was 2 m/s. The ground sampling distance (GSD) is 0.27 cm/pixel. The actual pixels of the camera are 20MP, the aperture size is F2.8 - F11, the autofocus range is from 1 m to infinity, and the shutter speed is set to 1/500s. The size of each image is 5472 × 3648 pixels, and the DPI of the images is 72 pixels/inch. UAV data collection was conducted weekly throughout the growing season.

### 2.3 Data Preprocessing

The images collected by the drone will be stitched into an orthomosaic by Agisoft Metashape (Agisoft LLC, St. Petersburg, Russia), in which geometric distortions have been corrected and the images have been color balanced. The excess portion of the orthomosaic is clipped and then further cropped into a small image size of 640×640 pixels to prevent exceeding the graphics card’s video memory during training.

### 2.4 Image Annotation

All labels in this study were generated by AnyLabeling. AnyLabeling combines the features of two labeling tools, LabelImg and Labelme. With the AI support of YOLO and Segment Anything Model (SAM) (Kirillov et al., 2023), it can help us semi-automate labeling. This study used the CPU version of AnyLabeling to label UAV images and used SAM for auxiliary labeling. All potato plants and weed outlines in the image are surrounded by polygons. In this study, with the assistance of SAM, 130 images in the training set were annotated, including 659 potatoes and 934 weeds, for a total of 1593 instances. There are 20 images in the validation set labeled, including 82 potatoes and 138 weeds, for a total of 220 images. The annotation file format generated by AnyLabeling is json and can be used to train the Mask R-CNN model, while the annotation file required for YOLO model training is in txt file format. The annotation files in json format are converted into YOLO format by Labelme’s python conversion script.

### 2.5 Data Augmentation

Due to the limited clarity of drone images, we used the Real-ESRGAN model to enhance our drone images and improve the annotation quality of the dataset. In this study, the resolution of the original image was increased by two times, but the size of the image was kept unchanged. In YOLOv8 we use the default data enhancement parameters.

### 2.6 Model Training

#### 2.6.1 Mask R-CNN

Mask R-CNN (Region-based Convolutional Neural Network) (He et al., 2017) is a deep learning framework for simultaneous object detection and instance segmentation. It identifies objects in an image, such as weeds and crops, and generates high-precision pixel-level masks for each object. Mask R-CNN’s main components include the backbone network (e.g., ResNet with FPN) for feature extraction, RPN for generating candidate regions, ROI Align for adjusting candidate regions to a fixed size, classification and bounding box regression head for categorizing and refining bounding boxes, and the mask prediction head for generating segmentation masks (He et al., 2017). This framework excels in object detection and instance segmentation, making it a powerful tool for various computer vision applications.

The Mask R-CNN model in this study was trained based on Detection2, the operating system is Ubuntu 22.04.4 LTS, and the GPU used is RTX3070Ti (Nvidia, California, United States). The first step is to install Anaconda and create a Python virtual environment for training the model. The Python version of the virtual environment used in this study is 3.8.18. Then install the packages and GPU environment required by Detection2 in the virtual environment, such as the 545.29.06 GPU driver, 11.8 version of CUDA, 8.7 version of CuDNN, 2.2.1 version of PyTorch, 1.24.3 version of numpy, 10.2.0 Version of Pillow, 0.17.1 version of torchvision, 4.9.0 version of cv2, etc. Due to compatibility issues, be very careful with the version of the installation package. The second step is to prepare the data set. We prepared a training set containing 130 images and a validation set of 20 images and converted the data set into the COCO format required by Detectron2. Each image has a corresponding annotation file, including a polygon mask and category labels. Before training, register the dataset into Detectron2. Then configure the parameters of the Mask R-CNN model, train and evaluate the model.

#### 2.6.2 YOLOv8

YOLOv8 (You Only Look Once version 8) represents a significant advancement in the field of object detection and segmentation, building upon the efficient, real-time performance characteristics of its predecessors. This framework integrates segmentation capabilities directly into the YOLO architecture, enabling simultaneous object detection and instance segmentation. YOLOv8 retains the core principle of the YOLO series: processing images in a single forward pass through the network, which ensures high-speed performance (Adarsh et al., 2020). However, it introduces several enhancements to improve accuracy and segmentation quality. YOLOv8 extends the traditional YOLO detection head to include a segmentation branch, which predicts segmentation masks along with bounding boxes and class probabilities (Terven et al., 2023). This allows for precise pixel-level delineation of objects within the image. The backbone network in YOLOv8, often based on advanced architectures like CSPDarknet (Bochkovskiy et al., 2020) or custom-designed variants, extracts richer and more detailed features from input images. This improves the model’s ability to identify and segment objects with greater accuracy. YOLOv8 employs specialized loss functions that balance the accuracy of object detection and the quality of segmentation masks, this ensures that both tasks are optimized simultaneously, leading to better overall performance (Sapkota et al., 2024). Despite its enhanced capabilities, YOLOv8 maintains the real-time processing speed characteristic of the YOLO family. This makes it suitable for applications requiring fast and accurate segmentation, such as autonomous driving, video surveillance, and precision agriculture (Terven et al., 2023).

The operating system for training the YOLOv8 model is Windows 11, the GPU is 3070Ti, the driver version is 552.22, the CUDA version is 12.3, the python version is 3.11, and the PyTorch version is 2.2.1. The first step is to use Anaconda to create a virtual environment required for training the model and install the dependencies and libraries for YOLOv8. The second step involves preparing the dataset like Mask R-CNN, but with labels in the YOLO format. In this format, class represents the object class index (starting from 0), and x_center, y_center, width, and height repr4esent the object’s center coordinates and dimensions. These values are normalized relative to the image size (ranging from 0 to 1). The third step creates a configuration file to specify the location of the data set and other parameters. Furthermore, we use the pre-trained YOLOv8 model as a base and then fine-tune it. Finally, the trained model is used for inference and evaluation.

### 2.7 Model Evaluation

The evaluation of the model is based on the mean average precision (mAP) metric, mAP is a metric used to evaluate the performance of object detection models. It comprehensively considers the accuracy and precision of the model in detecting objects and is widely used in various detection tasks in the field of computer vision, such as target detection and instance segmentation. The mAP is calculated as the average of the precision-recall curve, where the precision is the ratio of the number of true positive predictions to the total number of predictions, and the recall is the ratio of the number of true positive predictions to the total number of ground truth objects. The Precision-Recall curve refers to the curve obtained by plotting Precision and Recall under different thresholds. The horizontal axis is Recall, and the vertical axis is Precision. AP is obtained by calculating the area under the Precision-Recall curve. The mAP is the average of the AP values of all classes. The formula for calculating Precision, Recall, AP, and mAP is as follows:

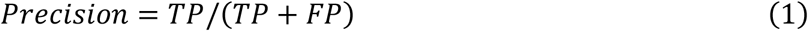

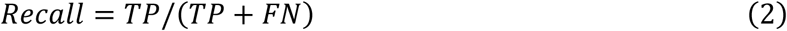

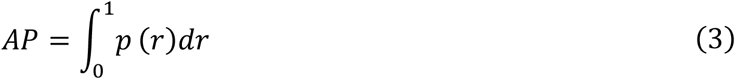

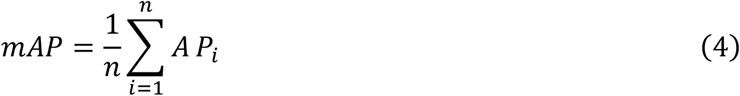

Where TP is the number of true positive predictions, FP is the number of false positive predictions, FN is the number of false negative predictions, p(r) is the precision value at recall r, n is the number of classes, and APi is the average precision of class i.

In addition, semantic segmentation is commonly evaluated using the Intersection over Union (IoU) metric. IoU is calculated as the ratio of the intersection area of the predicted mask and the ground truth mask to the union area of the two masks. The formula for calculating IoU is as follows:

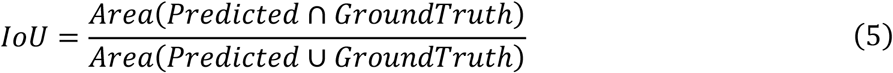

Mean IoU (mIoU) is the average of the IoU values of all classes. The formula for calculating mIoU is as follows:

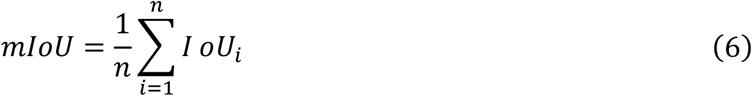

Where n is the number of classes, IoUi is the IoU of class i.

Pixel Accuracy (PA) is another metric used to evaluate semantic segmentation models, which calculates the ratio of the number of correctly classified pixels to the total number of pixels in the image. The formula for calculating Pixel Accuracy is as follows:

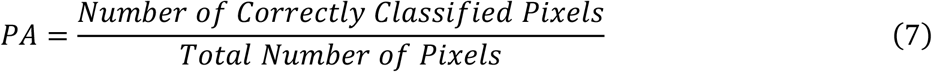

Class Pixel Accuracy (CPA) is the ratio of the number of correctly classified pixels of a specific class to the total number of pixels of that class. The formula for calculating CPA is as follows:

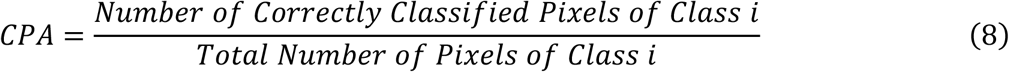

Mean Pixel Accuracy (MPA) is the average of the Pixel Accuracy values of all classes. The formula for calculating MPA and MCA is as follows:

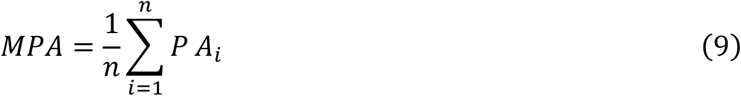

Where n is the number of classes, PAi is the Pixel Accuracy of class i. Dice Coefficient (DC) is another metric used to evaluate semantic segmentation models, which calculates the similarity between the predicted mask and the ground truth mask. The formula for calculating Dice Coefficient is as follows:

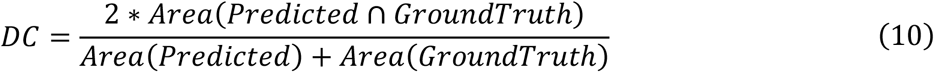

### 2.8 Model Application

This study uses a trained crop-weed segmentation model to infer images of field plots. By applying the shoelace formula to calculate the area of the predicted masks, the crop coverage rate and weed coverage rate for each plot are determined. The research further investigates the impact of weed and nitrogen treatments on potato yield.

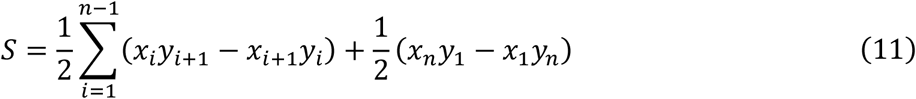

Where S is the area of the polygon, n is the number of vertices, and (x1,y1),(x2,y2),…,(xn,yn) are the coordinates of the vertices.

This study then used analysis of variance (ANOVA) and linear regression analysis to explore the effects of weed treatment, nitrogen fertilizer treatment on weed coverage, potato canopy coverage and final yield.

## 3 Results

### 3.1 UAV Images and Annotation

Figure 1 shows the orthomosaic of the field. As shown in Figure 2, the barrel distortion caused by the wide-angle lens is eliminated by Agisoft Metashape. In Metashape, lens distortion from drone imagery is corrected through the use of a camera calibration model that identifies and rectifies radial and tangential distortions. The process involves automatic image matching and alignment, application of geometric corrections, and orthophoto projection to map the corrected images onto a planar coordinate system. This methodology effectively eliminates lens distortions, resulting in high-precision orthophotos. Figure 3 shows the labeling results generated by SAM.

**Figure 1:**
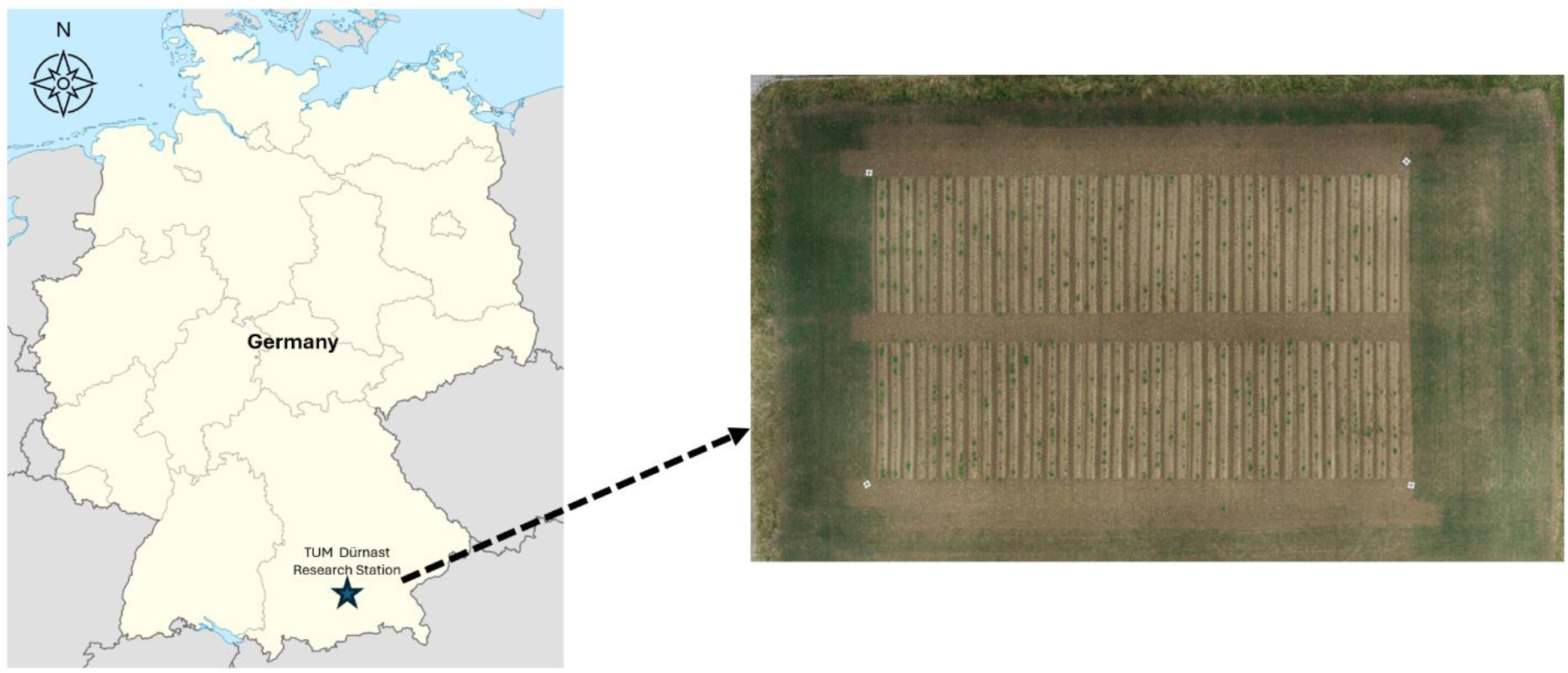
Experiment site and field othomosaic

**Figure 2:**
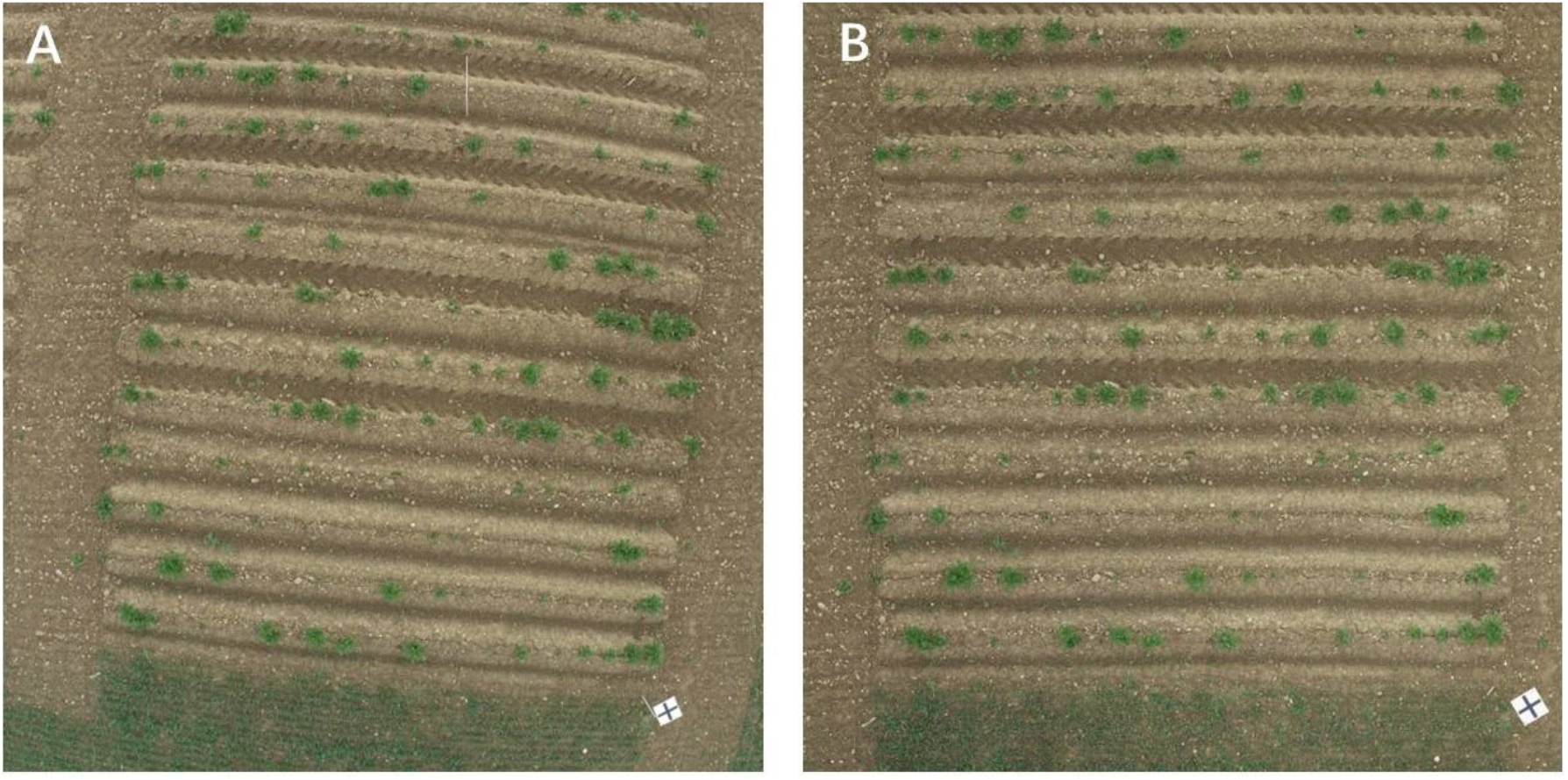
Distortion correction. A is the original image, and B is the distortion-corrected image.

**Figure 3:**
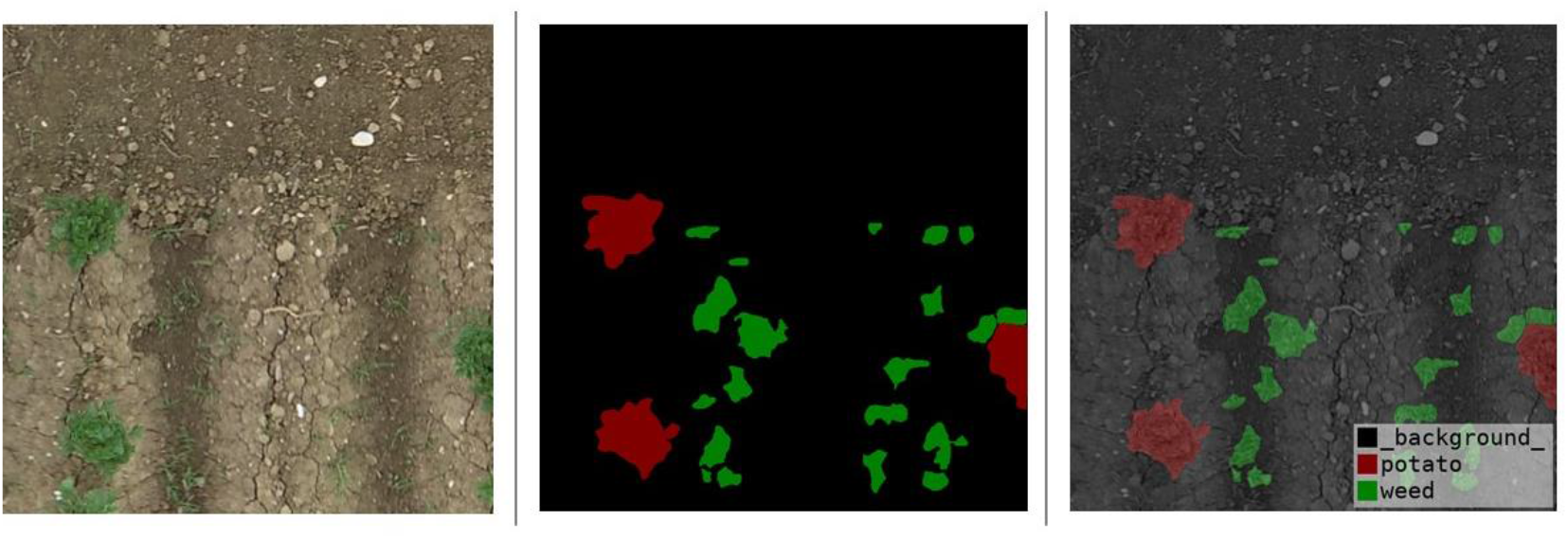
Annotation visualization.

We evaluated the average annotation speed for 10 instances manually and with SAM assistance. Manual annotation averaged 5.84 seconds per instance, whereas SAM-assisted annotation averaged 19.2 seconds per instance. This demonstrates that SAM-assisted annotation is 3.3 times more efficient than manual annotation

### 3.2 Data Augmentation and Super-resolution Reconstruction

Figure 4 illustrates the data enhancement techniques used by YOLOv8, simulating images under various environments. These techniques include Mosaic, RandomAffine, MixUp, Image Blur, Transform, HSV Color Space Enhancement, Random Horizontal Flip. The primary purpose of data augmentation is to artificially increase the diversity of the training dataset, thereby improving the model’s robustness to variations in object appearance, orientation, and environmental conditions. By exposing the model to a wider range of scenarios, data augmentation helps in reducing overfitting and improving the model’s ability to generalize to new, unseen data.

**Figure 4:**
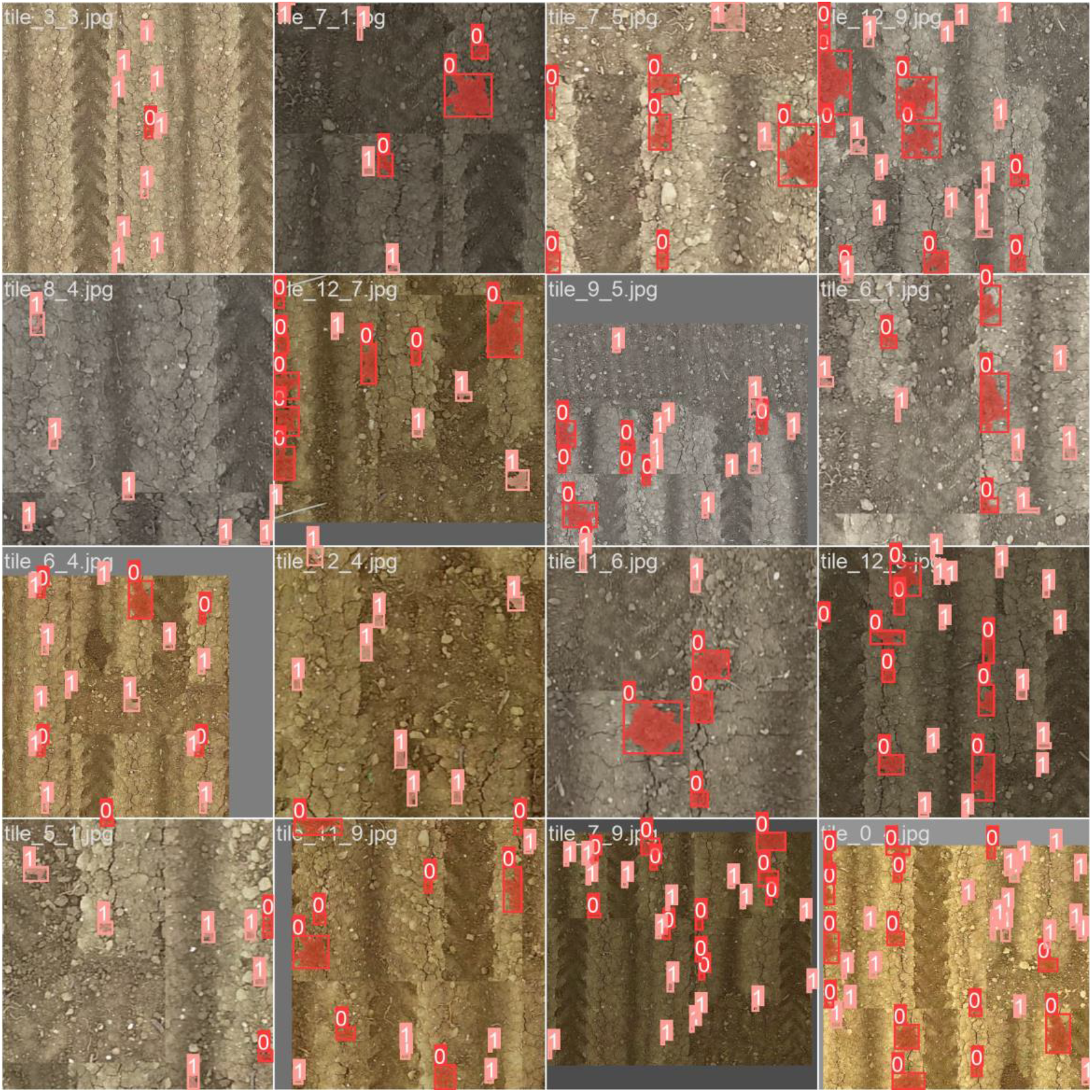
Data augmentation example. Includes Mosaic, RandomAffine, MixUp, Image Blur, Transform, HSV Color Space Enhancement, Random Horizontal Flip.

Figure 5 presents the UAV images before and after Real-ESRGAN super-resolution reconstruction. The original resolution in Figure 5A is 640×640 pixels, while Figure 5B, after 2x super-resolution reconstruction, has a resolution of 1280×1280 pixels, resulting in clearer object edges.

**Figure 5:**
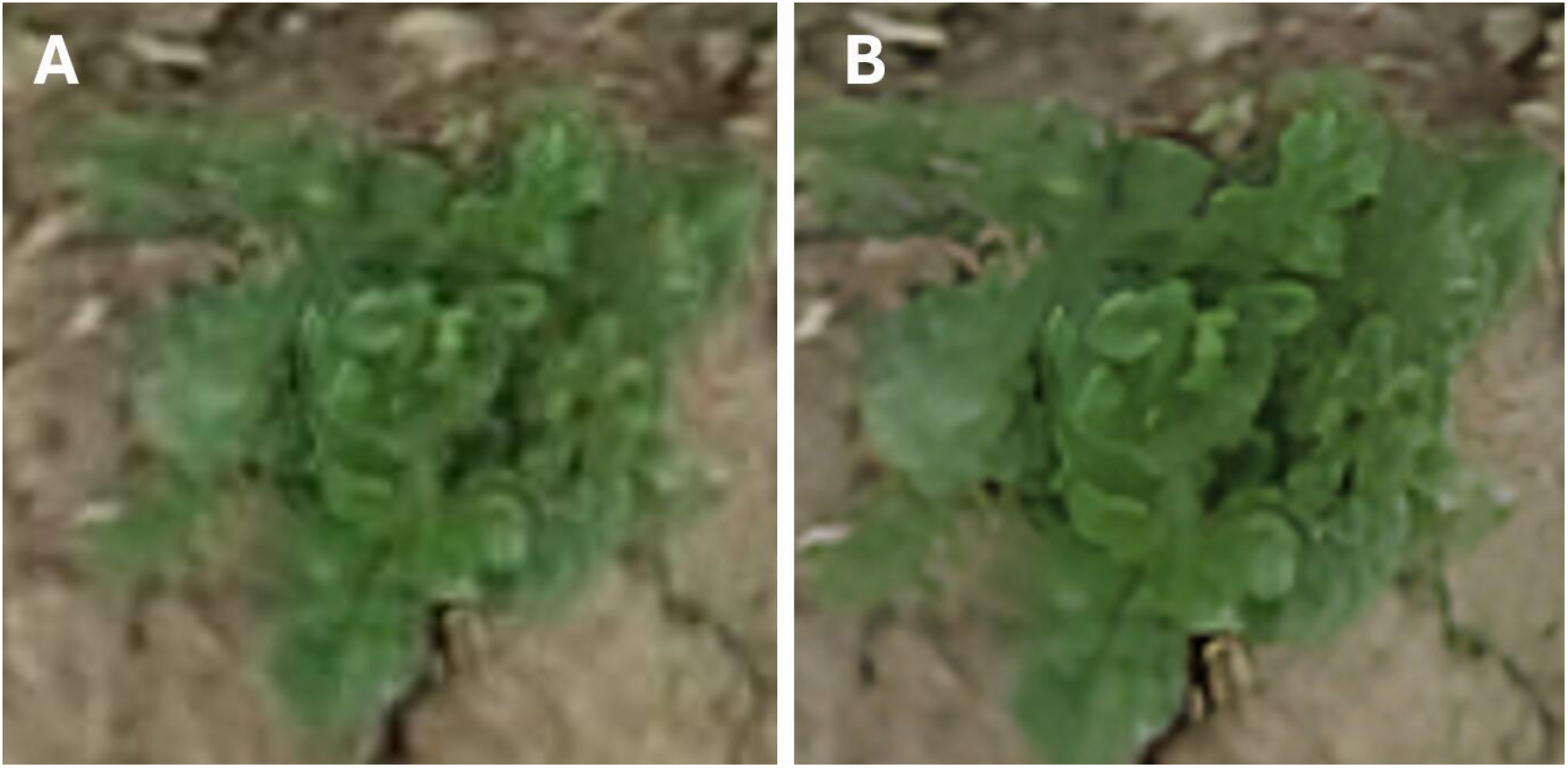
Real-ESRGAN super-resolution reconstruction. A is the original image, and B is the super-resolution reconstructed image.

### 3.3 Training results and inference performance of Mask R-CNN and YOLOv8

#### 3.3.1 Mask R-CNN

Figure 6 illustrates the indicator trends of the Mask R-CNN model. With the total loss gradually decreases as training progresses (Figure 6A), indicating continuous learning and optimization, with a general downward trend despite some fluctuations. Figure 6B shows that the model’s accuracy improves with the number of iterations, reflecting enhanced detection and segmentation capabilities. The upward curve indicates increasing accuracy and reliability.

**Figure 6:**
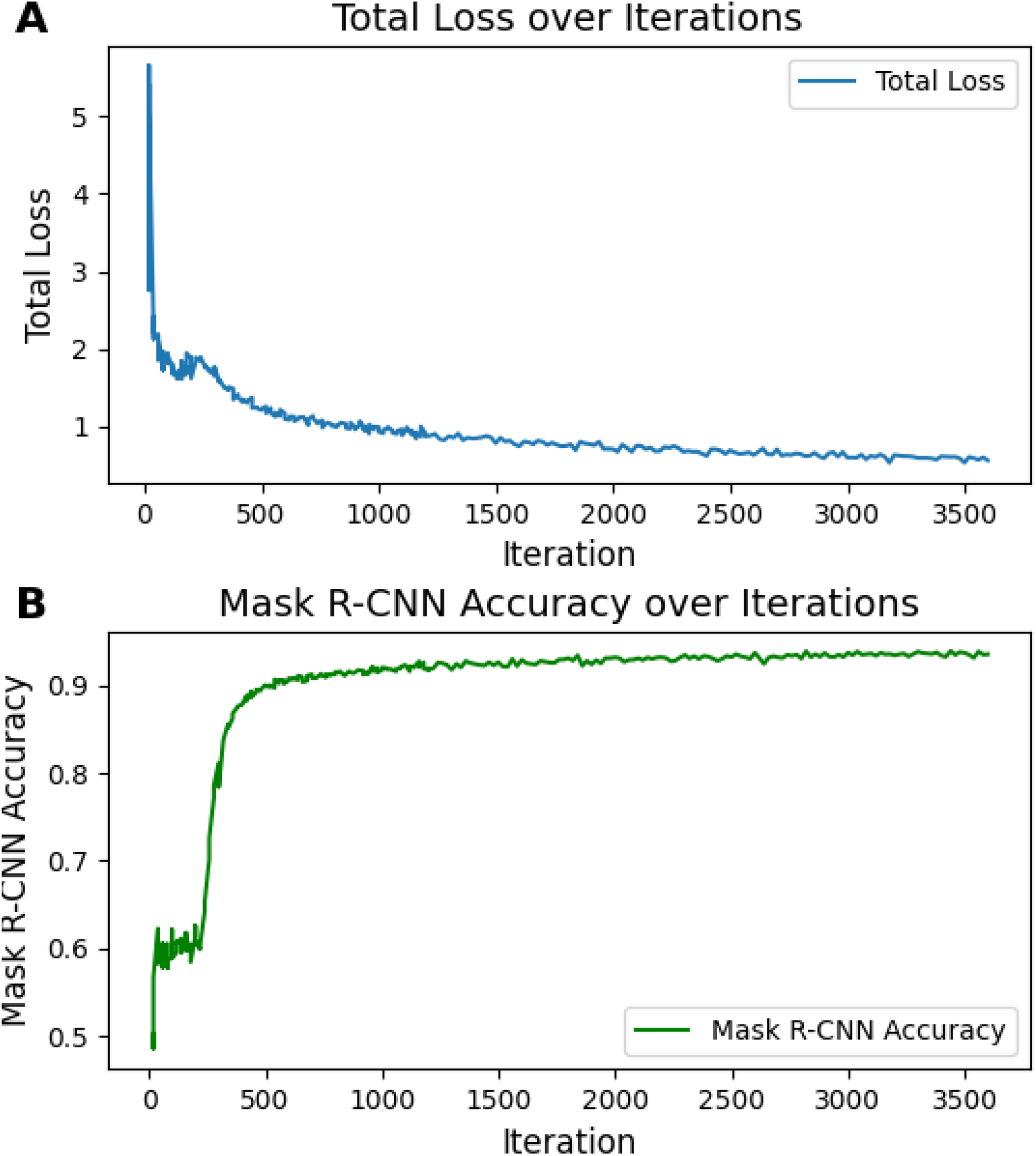
Mask R-CNN results. A is the trend graph of total loss over iterations, B is the trend graph of accuracy over iterations.

Figure 7 presents the Average Precision (AP) results for the Mask R-CNN model, while Table 1 details the corresponding AP indicators. Initially, with a smaller number of iterations (e.g., the first 200), the AP values are relatively low, indicating that the model’s performance has not yet been fully realized. As the iterations increase into the hundreds, the AP values rise significantly, reflecting the model’s improvement through continuous learning and optimization. Beyond 1000 iterations, most AP values stabilize, and the model’s performance plateaus at a high level, with the rate of improvement slowing down. Figure 8 demonstrates the inference results of the Mask R-CNN model, which accurately detects and segments potatoes and weeds, producing clear and precise segmentation outcomes.

**Table 1:** Mask R-CNN AP indicators.

| Indicators | Description |
| --- | --- |
| bbox/AP | Average accuracy of bounding boxes. |
| segm/AP | average accuracy of segmentation. |
| bbox/AP-potato | Average accuracy for bounding box detection of potatoes. |
| bbox/AP-weed | Average accuracy of bounding box detection of weeds. |
| segm/AP-potato | Average accuracy for segmenting and detecting potatoes. |
| segm/AP-weed | Average accuracy of segmentation detection of weeds. |
| bbox/AP50 | The average accuracy of the bounding box when the intersection-over-union ratio (IoU) is 50%. |
| bbox/AP75 | The average accuracy of the bounding box when the intersection and union ratio is 75%. |
| segm/AP50 | The average segmentation accuracy when the intersection and union ratio is 50%. |
| segm/AP75 | The average segmentation accuracy when the intersection and union ratio is 75%. |

**Figure 7:**
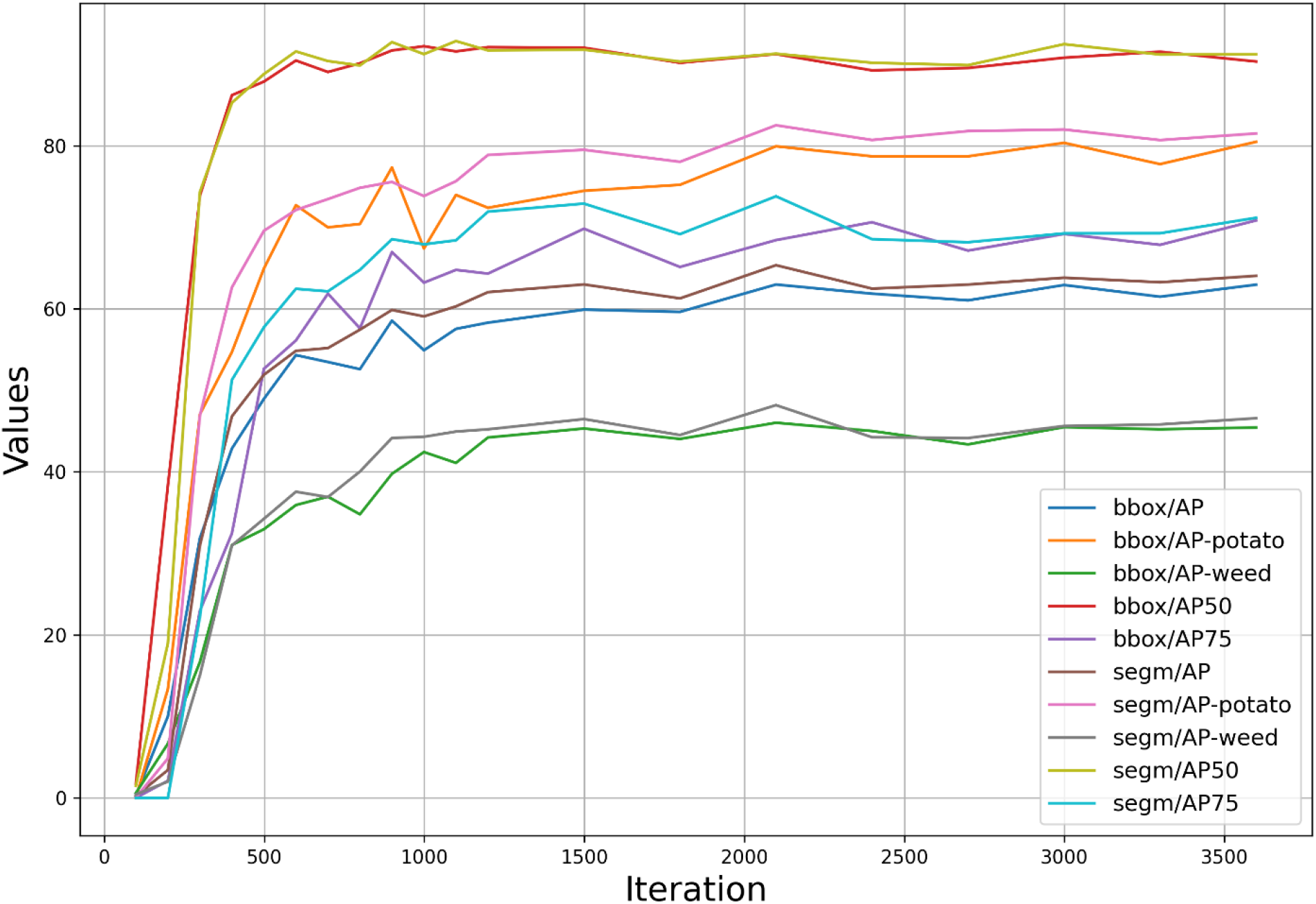
Mask R-CNN AP results.

**Figure 8:**
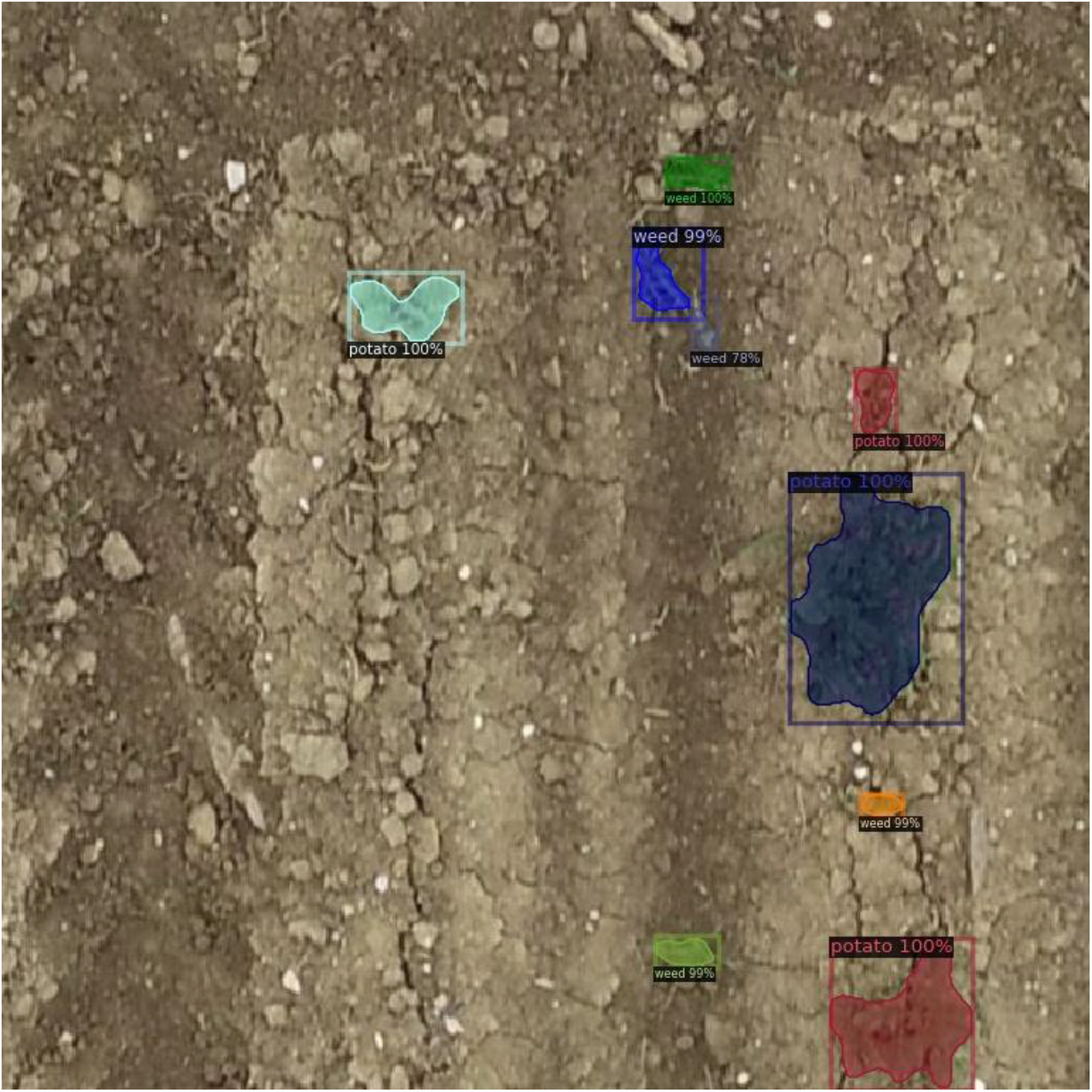
Mask R-CNN Inference example.

#### 3.3.2 YOLOv8

Figure 9 illustrates the indicator trend changes of the YOLOv8 models. These figures display the overall trends of the loss function and evaluation metrics during training and validation. The curves reveal performance improvements across different iterations. Generally, the loss function curve exhibits a downward trend, while the evaluation metrics show an upward trend, indicating continuous optimization and performance enhancement.

**Figure 9:**
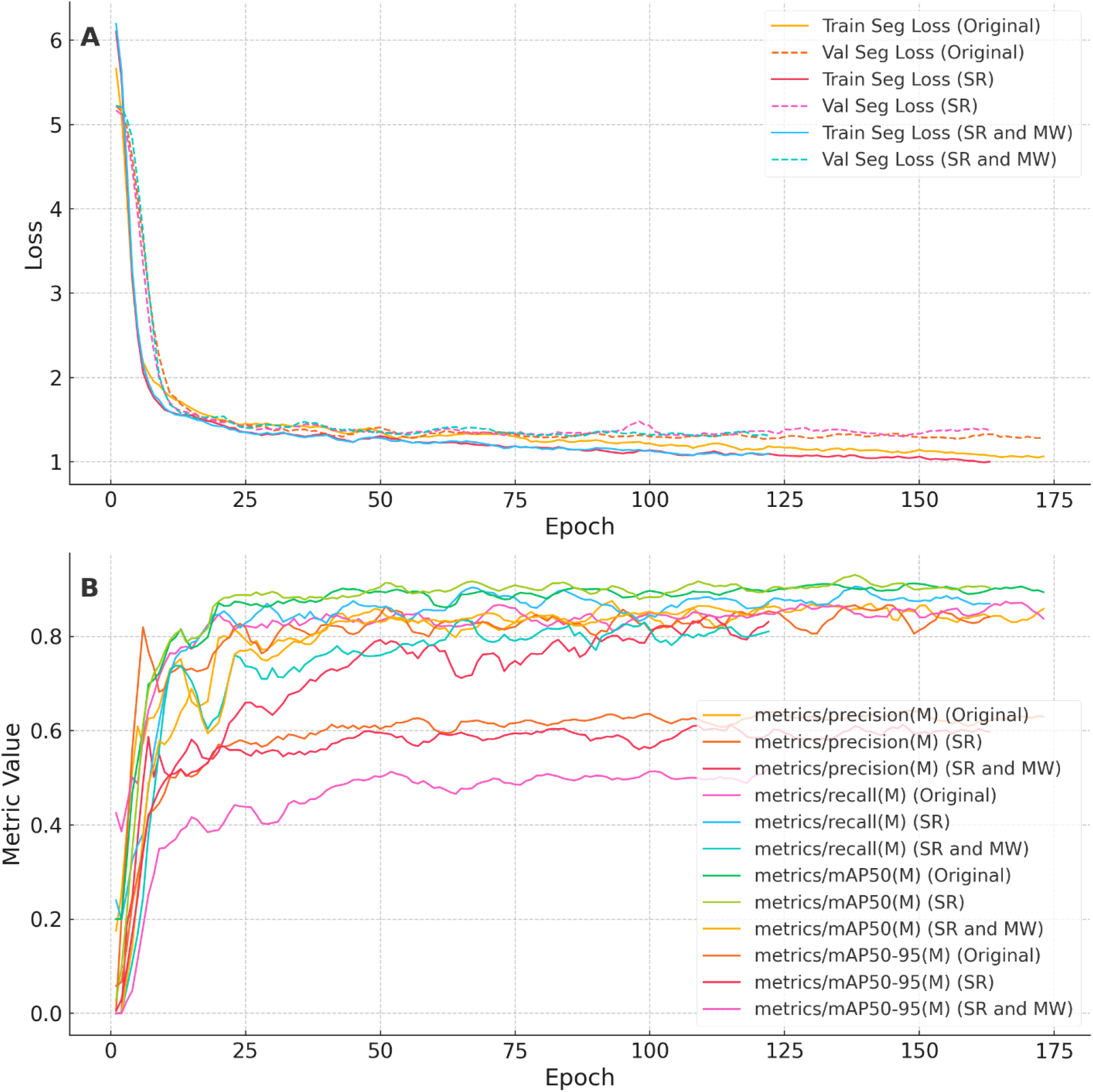
YOLOv8 results of Original, Super-resolution reconstructed (SR), and Super-resolution reconstructed and multiple weed categories (SR and MW). A. Training and Validation loss, B. Accuracy metrics

Train/seg_loss: Shows the trend of segmentation loss, with the curve stabilizing from high to low, indicating segmentation task optimization.

Val/seg_loss: Reflects the segmentation loss trend on the validation set, decreasing and tending to stabilize.

Metrics/precision (M): Shows the trend of segmentation accuracy, increasing with iterations and stabilizing.

Metrics/recall (M): Displays the trend of segmentation recall, also increasing with iterations and stabilizing.

Metrics/mAP50 (M): Demonstrates the change in mean average precision (mAP) at an IoU threshold of 50%, with curves rising rapidly and stabilizing.

Metrics/mAP50-95 (M): Indicates the change in mean average precision (mAP) at multiple IoU thresholds (50%-95%), with curves rising rapidly and stabilizing.

These trends collectively indicate the model’s consistent optimization and improved performance across training and validation phases.

Figure 10 A (1) shows the precision-recall of potatoes, which shows that the precision of potatoes was very high, reached 0.983, the precision-recall of weeds was 0.821, which was lower than that of potatoes. The average precision (mAP@0.5) of all categories is 0.902.

**Figure 10:**
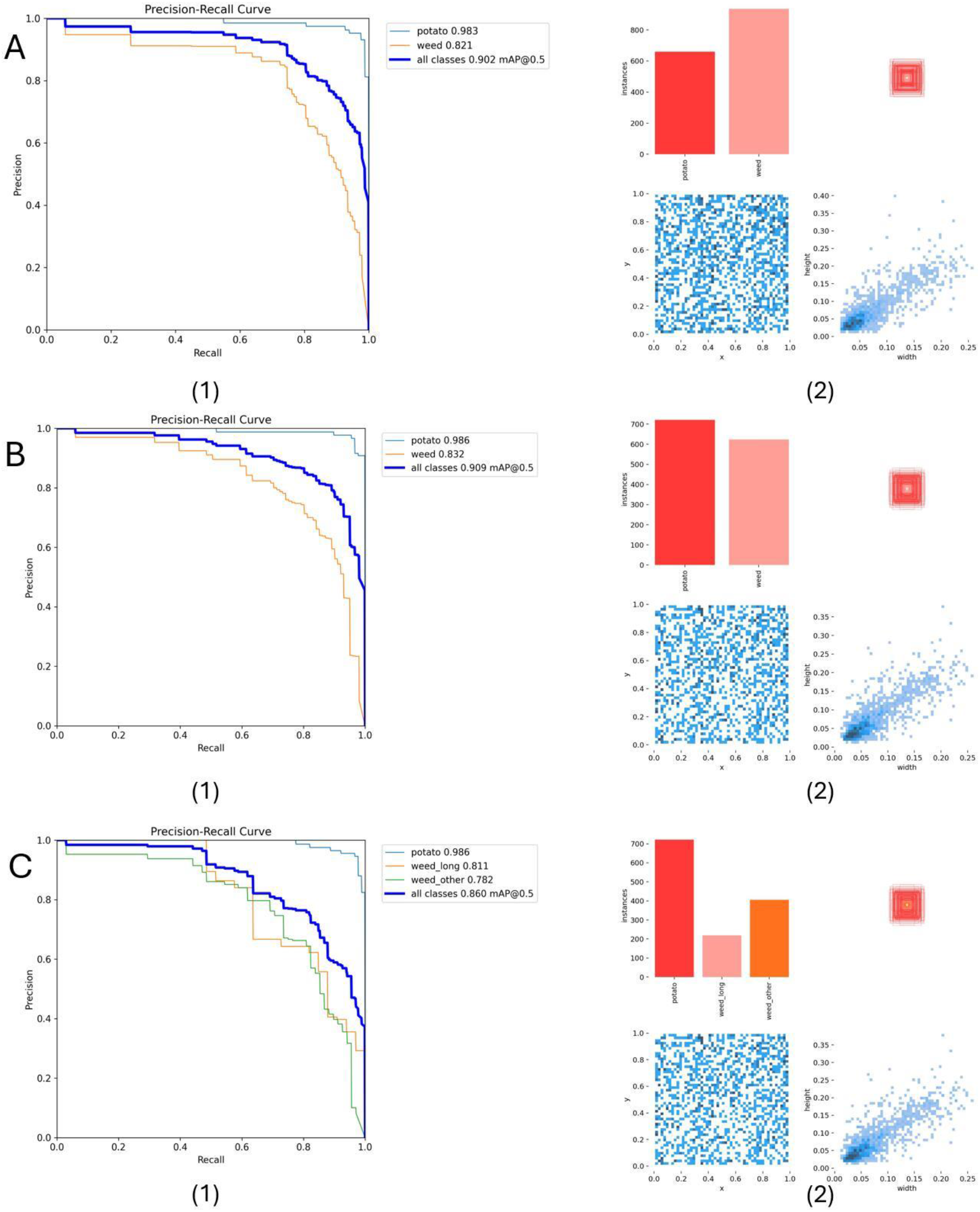
YOLOv8 PR and labels results. A. Original, B. Super-resolution reconstructed, C. Super-resolution reconstructed and multiple weed categories. (1) is the PR curve of the model. (2) is the label results of the model.

Figure 10 B (1) shows the precision-recall of potatoes is 0.986. The precision-recall of weeds is 0.832, which is higher than the precision of weeds in Part A. The average precision (mAP@0.5) of all categories, which is 0.909, slightly higher than Part A.

Figure 10 C (1) shows the precision-recall of potatoes is 0.986, which is consistent with Part B. The precision-recall of long weeds is 0.811. The precision-recall of other weeds is 0.782. The average precision (mAP@0.5) of all categories is 0.860, lower than A and B. Figure 10 C (2) shows that the three models detect about the similar number of potato instances, but model A detects significantly more weed instances.

Figure 11 shows the segmentation results of the YOLOv8 model. Original Model: PA: 0.9989, CPA: 0.5271, MPA: 0.5150, MIOU: 0.5150, Mean Dice: 0.5298. The original model shows a high PA but lower performance in CPA, MPA, MIOU, and Mean Dice compared to PA. Super-resolution (SR) Model: PA: 0.9990, CPA: 0.5250, MPA: 0.5116, MIOU: 0.5116, Mean Dice: 0.5234. This model has slightly improved PA but a small decline in CPA, MPA, MIOU, and Mean Dice compared to the original model. Super-resolution with Multiple Weed Categories (SR and MW) Model: PA: 0.9985, CPA: 0.5122, MPA: 0.5089, MIOU: 0.5089, Mean Dice: 0.5185, This model shows a slight reduction in PA, CPA, MPA, MIOU, and Mean Dice compared to the other two models. Pixel Accuracy (PA) is very high across all models, indicating that most pixels are correctly classified. Class Pixel Accuracy (CPA), Mean Pixel Accuracy (MPA), Mean Intersection over Union (MIOU), and Mean Dice show more variability, indicating that while pixel classification is generally accurate, the detailed performance across different classes and segmentation overlap can vary. The Super-resolution (SR) model has a slight advantage in PA but doesn’t significantly improve other metrics compared to the original model. The SR and MW model shows a slight decrease in performance across all metrics, suggesting that incorporating multiple weed categories may introduce complexity that affects overall segmentation accuracy.

**Figure 11:**
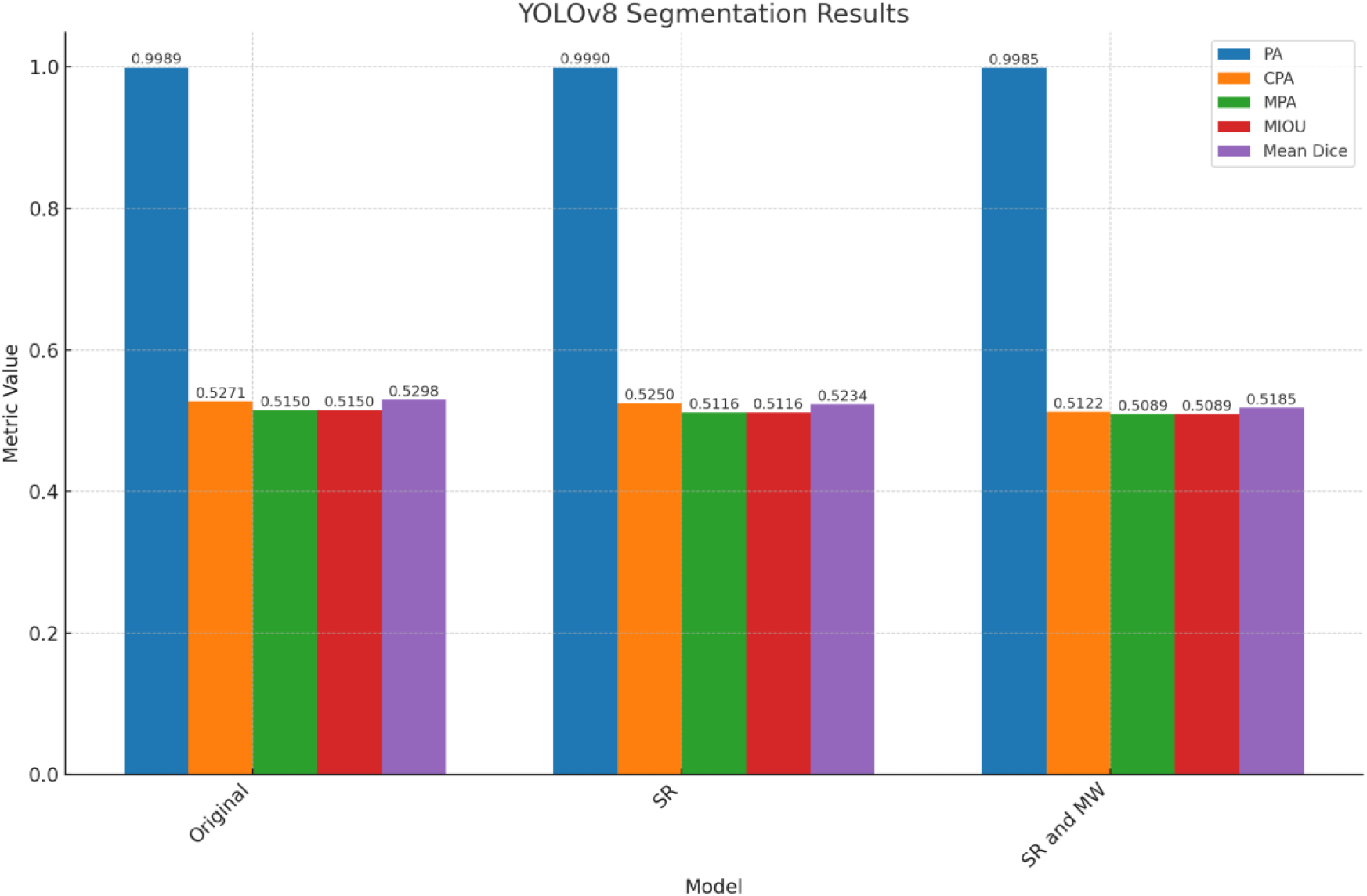
YOLOv8 segmentation results of original model, super-resolution reconstructed model (SR), and super-resolution reconstructed and multiple weed categories model (SR and MW).

Figure 12 shows the inference results of the YOLOv8 models. The models can accurately detect and segment potatoes and weeds in the image, and the segmentation results are clear and accurate. The models can also detect multiple weed categories, which can help farmers identify different types of weeds and take targeted measures to control them.

**Figure 12:**
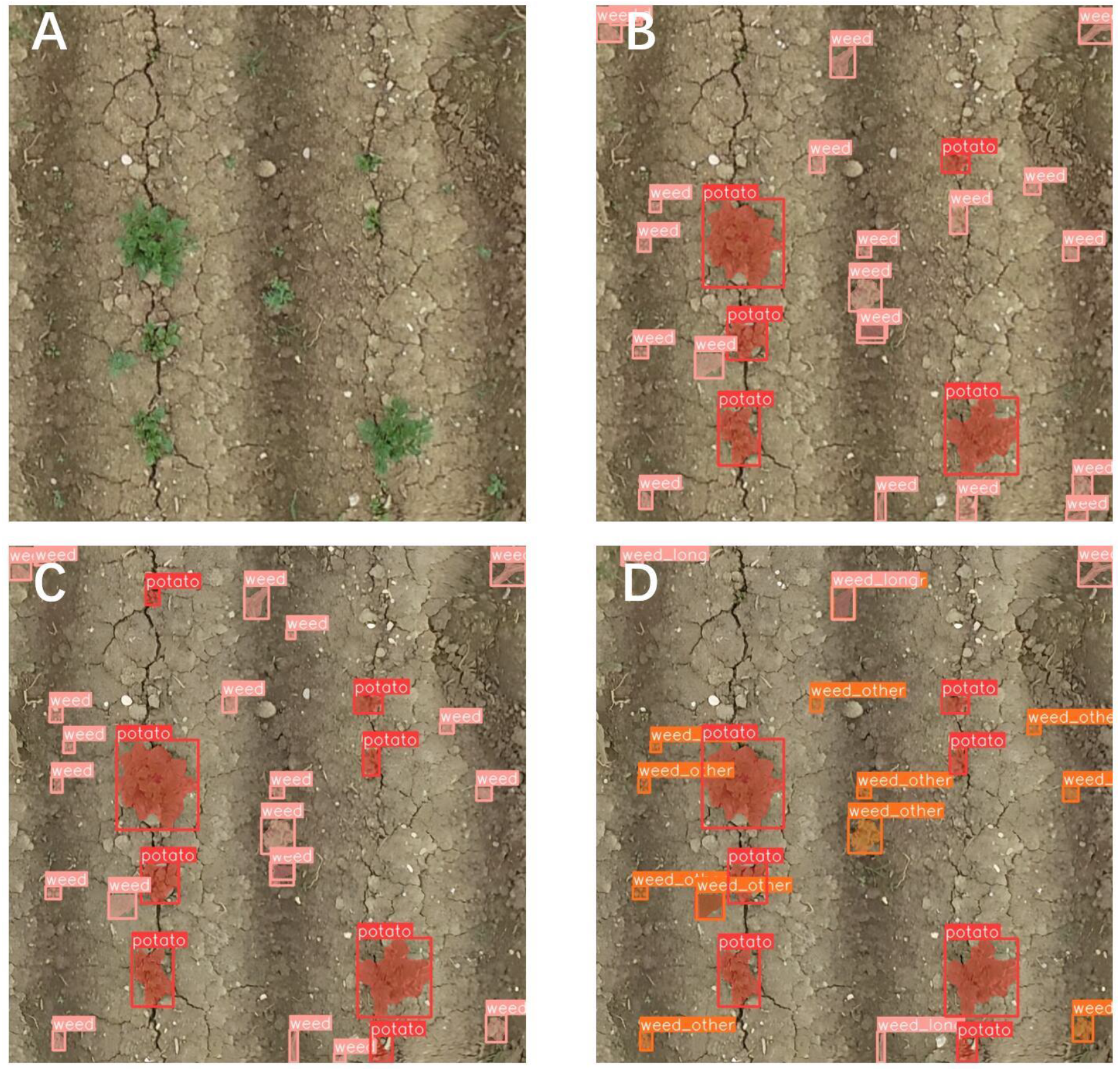
YOLOv8 Inference example. A. Raw image, B. Original, C. Super-resolution reconstructed, D. Super-resolution reconstructed and multiple weed categories.

### 3.4 Application of the YOLO Model

Figure 13 shows the original YOLO model used to segment the potato and weed coverage area of each plot. Table 2 is a summary of the field experiment data, including plot number, weed treatment, nitrogen fertilizer treatment, potato coverage area, weed coverage area, and potato yield.

**Table 2:** Overview of the field experiment results.

| Plot | Weed control | Nitrogen(kg N/ha) | Potato area(pixels) | Weed area(pixels) | Yield(dt(100kg)/ha) |
| --- | --- | --- | --- | --- | --- |
| 1 | control | 150 | 177768 | 50526 | 393.889 |
| 2 | chemical | 150 | 336231 | 27083 | 452 |
| 3 | mechanical | 150 | 263423 | 16239 | 514.556 |
| 4 | mechanical | 150 | 193646 | 9700 | 470.778 |
| 5 | chemical | 150 | 124780 | 16728 | 475 |
| 6 | control | 150 | 164923 | 42855 | 379.778 |
| 7 | chemical | 150 | 114664 | 12414 | 432.667 |
| 8 | mechanical | 150 | 267976 | 7870 | 415.333 |
| 9 | control | 150 | 133814 | 40910 | 396.667 |
| 10 | mechanical | 150 | 172552 | 31091 | 400.333 |
| 11 | control | 75 | 199114 | 109455 | 308.111 |
| 12 | chemical | 75 | 158497 | 29433 | 422.667 |
| 13 | mechanical | 75 | 127801 | 16753 | 436.556 |
| 14 | mechanical | 75 | 137742 | 16546 | 463 |
| 15 | chemical | 75 | 155804 | 20801 | 393.333 |
| 16 | control | 75 | 104856 | 21742 | 314.333 |
| 17 | chemical | 75 | 88812 | 18659 | 434 |
| 18 | mechanical | 75 | 97838 | 8106 | 383 |
| 19 | control | 75 | 88116 | 24061 | 384.333 |
| 20 | mechanical | 75 | 103884 | 18199 | 502.444 |

**Figure 13:**
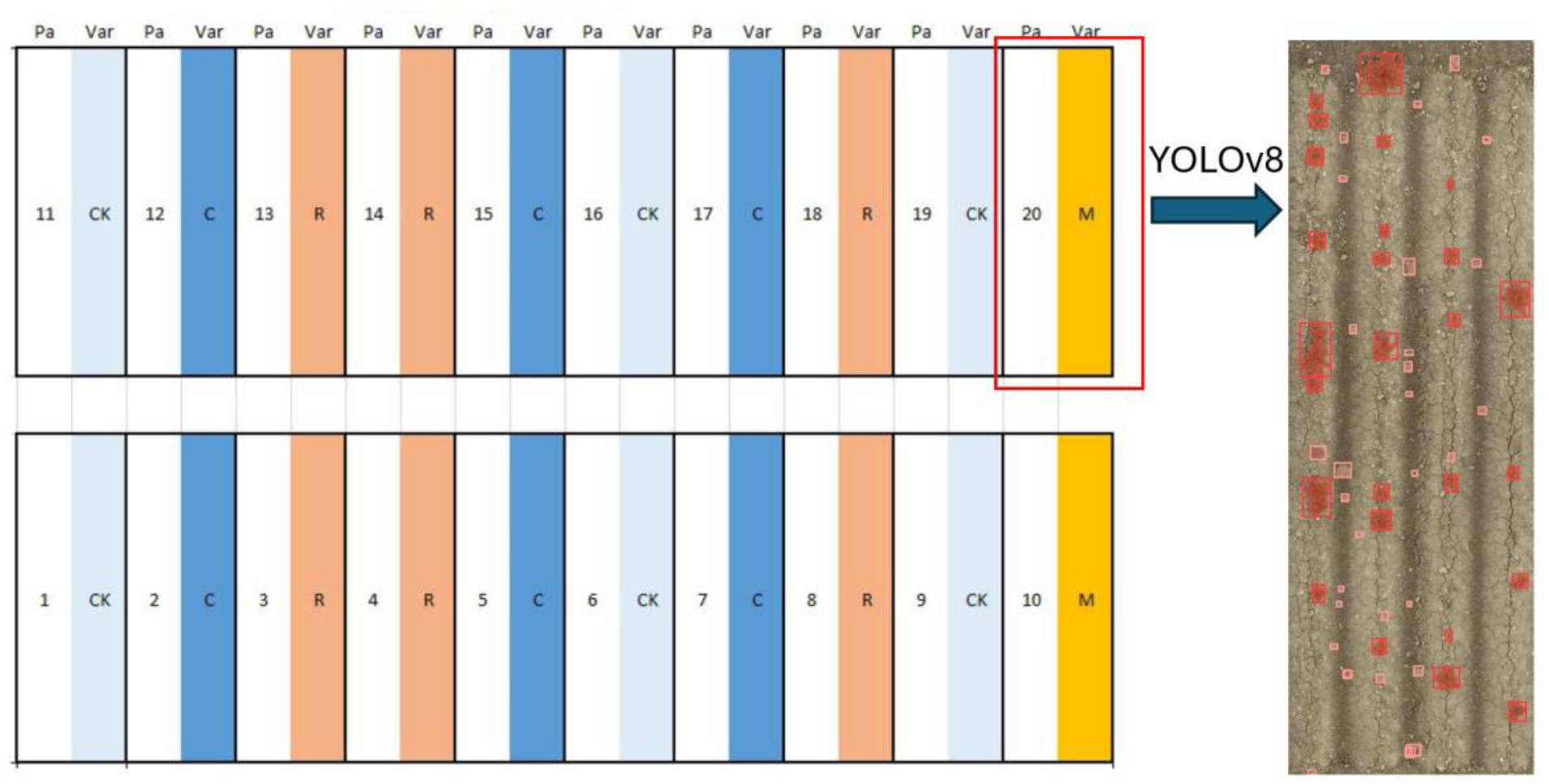
YOLOv8 application. YOLOv8 model extracts the coverage area of potatoes and weeds in each plot.

#### 3.4.1 Effects of Weed Control and Nitrogen Level on Yield

Figure 14 shows the potato yield under different weed control and nitrogen levels. The yield of potatoes under different treatments is different. The yield of potatoes under mechanical weed control is higher than that under chemical weed control and the control group with 75 kg N/ha. The yield of potatoes under chemical weed control is higher than that under mechanical and the control group with 150 kg N/ha.

**Figure 14:**
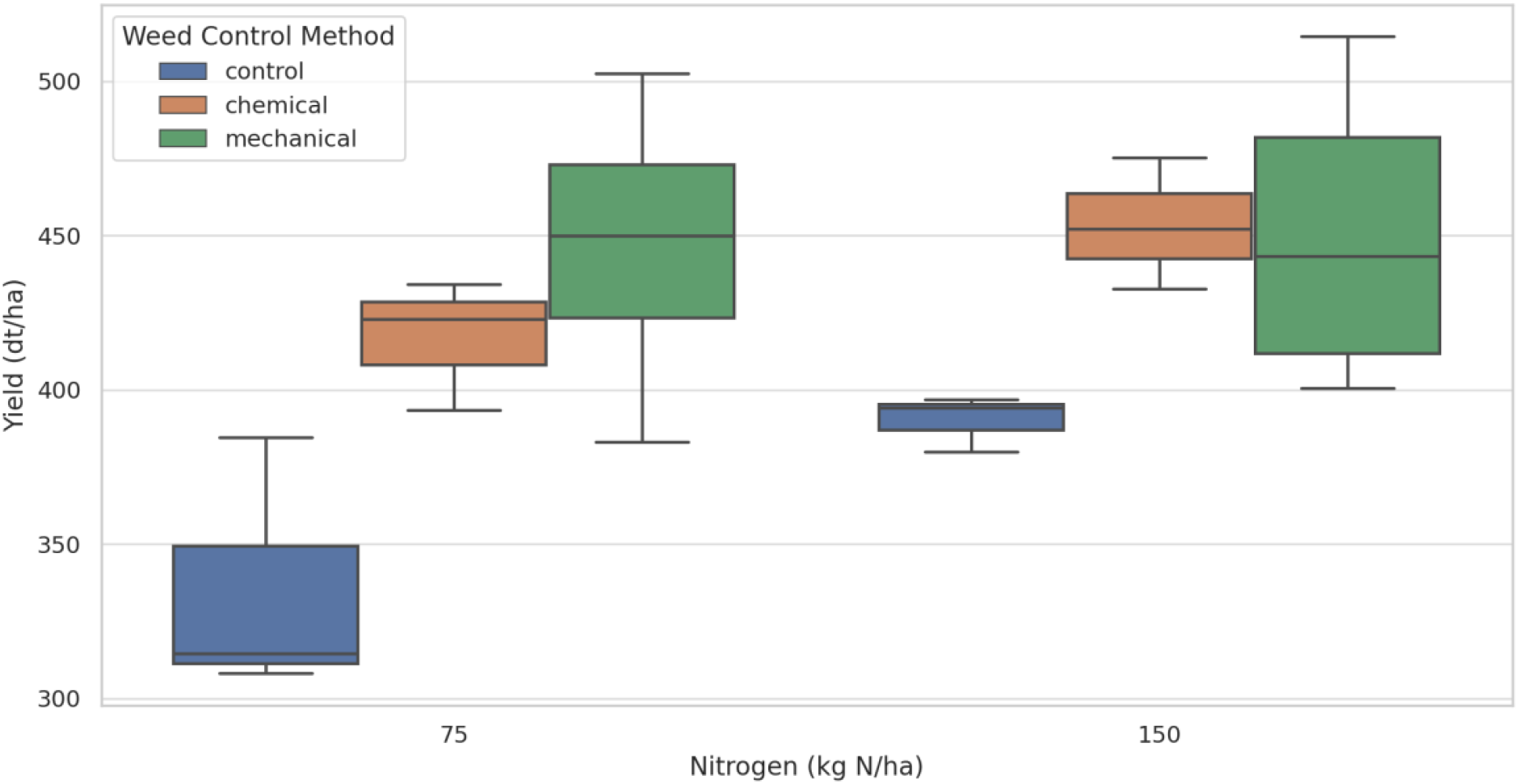
Potato yield under different treatments.

Table 3 shows the results of the ANOVA with the interaction between weed control and nitrogen. Weed control had a significant effect on the yield of potatoes, while the nitrogen level had no significant effect on the yield of potatoes. The interaction between weed control and nitrogen has no significant effect on the yield of potatoes. This means that their effects on yield appear to be independent, rather than interacting.

**Table 3:** ANOVA with interaction between Weed and Nitrogen.

|  | sum_sq | df | F | PR(>F) |
| --- | --- | --- | --- | --- |
| C(Weed_control) | 27282.954337 | 2.0 | 8.955096 | 0.003129 |
| C(Nitrogen) | 4182.526109 | 1.0 | 2.745665 | 0.119750 |
| C(Weed_control):C(Nitrogen) | 2312.431081 | 2.0 | 0.759010 | 0.486454 |
| Residual | 21326.480339 | 14.0 |  |  |

Interestingly, the yield of chemical treatment was significantly higher than control treatment (p = 0.0312, Table 4), the yield of mechanical treatment was also significantly higher than control treatment(p = 0.0033), but the yield difference between chemical treatment and mechanical treatment was not significant (p = 0.7177). Both chemical and mechanical treatments significantly increased potato yield compared with the control treatment, while the yield difference between chemical treatment and mechanical treatment was not significant.

**Table 4:** Turkey HSD test.

| group1 | group2 | meandiff | p-adj | lower | upper | reject |
| --- | --- | --- | --- | --- | --- | --- |
| chemical | control | 55.7162 | 0.0312 | 4.4562 | 106.9761 | True |
| chemical | mechanical | -16.5797 | 0.7177 | -67.8396 | 34.6802 | False |
| control | mechanical | -72.2959 | 0.0033 | -123.5559 | -21.0359 | True |

#### 3.4.2 Relationship between Potato and Weed Cover Area and Yield

Figure 15 shows the correlation matrix of the potato and weed coverage area and yield. There is a positive correlation between potato area and yield, meaning that the larger the potato planting area, the higher the yield. There is a negative correlation between weed area and yield, meaning that the larger the weed-infested area, the lower the yield. Table 5 and Figure 16 shows the results of the linear regression model with ‘Potato area’ and ‘Weed area’ as predictors for ‘Yield’. The R-squared of the model is 0.412, indicating that the model can explain about 41.2% of the variations in yield. The coefficient of Intercept is 421.1414, which represents the expected yield baseline in the absence of potato area and weed area. The coefficient of potato area is 0.0002, which means that for every additional pixel of potato plants, potato yield will increase by 0.02 kg/ha; while the coefficient of weed area is -0.0015, which means that for every additional weed pixel, yield will decrease by 0.15 kg/ha. Additionally, the model validity test yields an F statistic of 5.967 with an associated p-value of 0.0109, demonstrating that the model is statistically significant overall..

**Table 5:**
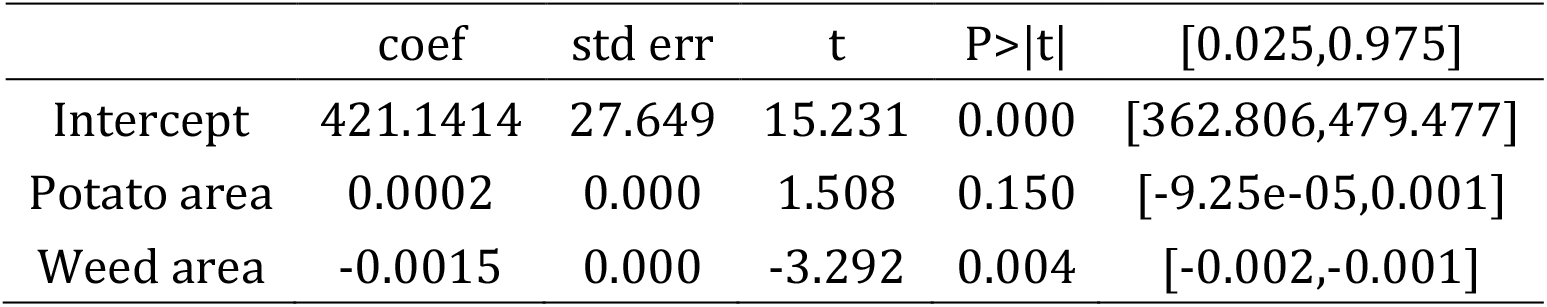
Linear regression model with ‘Potato area’ and ‘Weed area’ as predictors for ‘Yield’.

|  | coef | std err | t | P>\|t\| | [0.025,0.975] |
| --- | --- | --- | --- | --- | --- |
| Intercept | 421.1414 | 27.649 | 15.231 | 0.000 | [362.806,479.477] |
| Potato area | 0.0002 | 0.000 | 1.508 | 0.150 | [-9.25e-05,0.001] |
| Weed area | -0.0015 | 0.000 | -3.292 | 0.004 | [-0.002,-0.001] |

**Figure 15:**
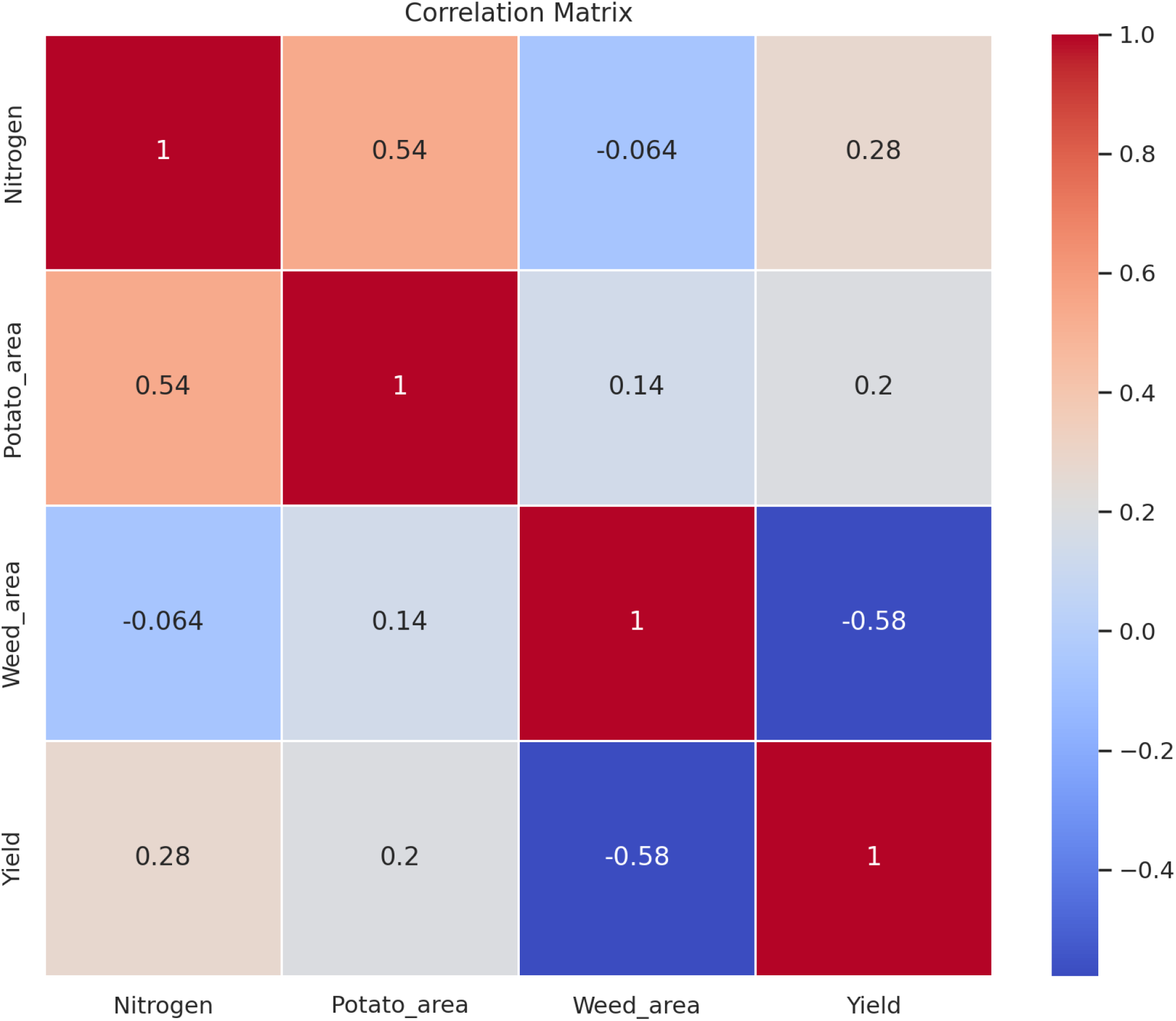
Correlation Matrix.

**Figure 16:**
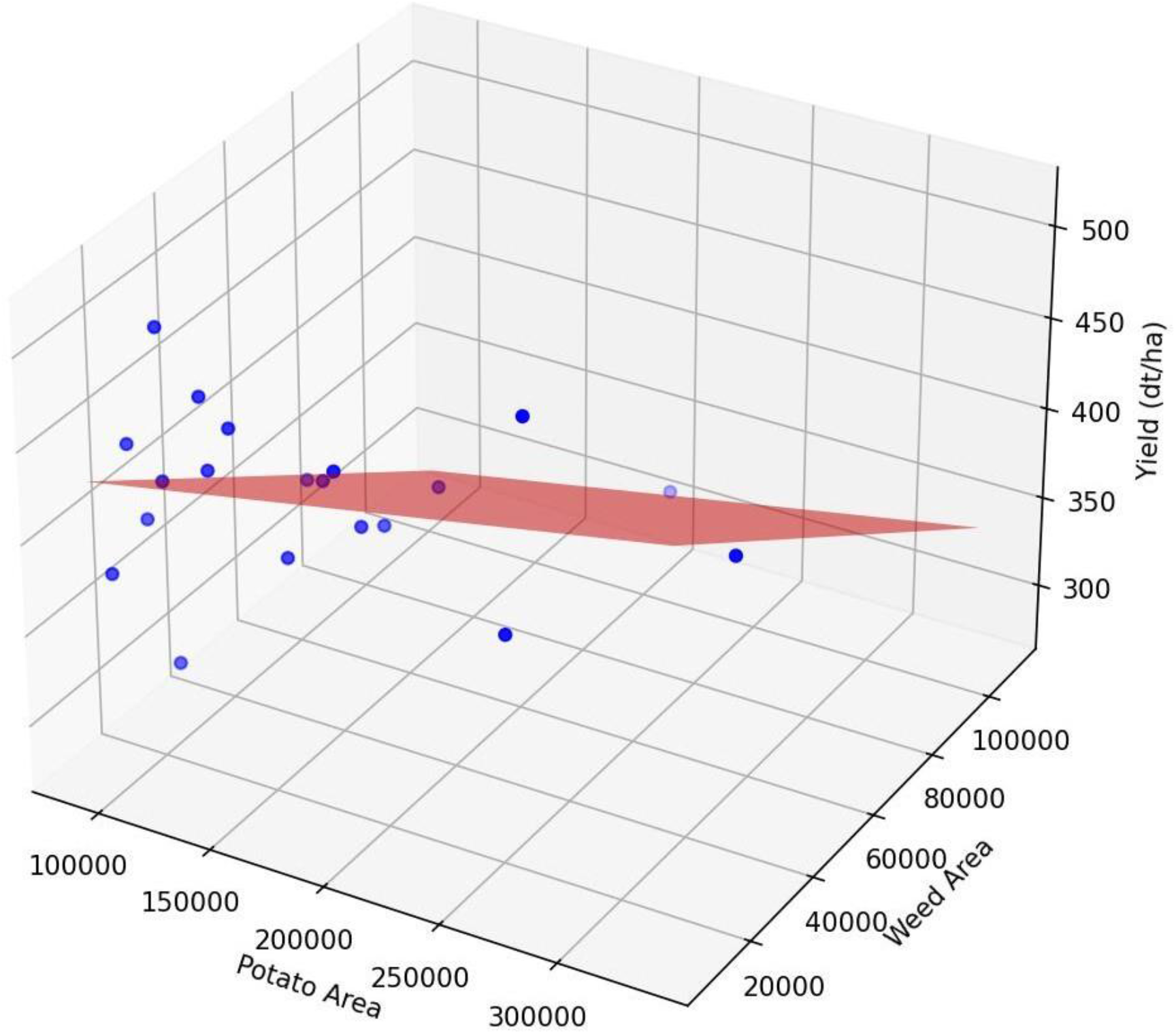
Multiple Linear Regression: Potato Area and Weed Area vs Yield

## 4 Discussion

### 4.1 Comparison between Mask R-CNN and YOLOv8

Kauer conducted a comprehensive comparison of various object detection and instance segmentation algorithms, including the YOLO series and Mask R-CNN, and YOLOv5 surpassed Mask R-CNN in terms of accuracy, speed, and computational efficiency for the segmentation task on the COCO dataset (Kaur and Singh, 2023).

In our study the accuracy of YOLOv8 and Mask R-CNN is very close, but YOLOv8 converges faster than Mask R-CNN, which can be attributed to several factors intricately related to the world of neural network architectures and training methods.

YOLOv8 is a more concise approach to object detection, using a single neural network to predict bounding boxes and class probabilities directly from the full image in a single evaluation (Hussain, 2023) This design is in stark contrast to the more complex structure of Mask R-CNN, which involves multiple stages and thus requires more computation per iteration (He et al., 2017). The world of single-shot detection methods, exemplified by YOLOv8, allows for faster processing times, and thus faster convergence.

The backbone network in YOLOv8 is generally redesigned to be more efficient and lightweight than the backbone network in Mask R-CNN (Hussain, 2023). This efficiency is critical in the rigors of training, where computational resources and time are paramount. By leveraging advanced feature extraction techniques and an optimized network architecture, YOLOv8 reduces the number of parameters and computational overhead, resulting in faster convergence.

In addition, the training strategy employed by YOLOv8 is carefully designed to outperform traditional methods. Techniques such as mosaic augmentation combine multiple images into one, providing ever-changing training data in each batch (Jocher et al. 2023). The model utilizes refined loss functions, such as IoU-based losses, to improve localization accuracy and stability (Su et al., 2024). Adaptive learning rate strategies, including dynamic adjustments, warm-up, and decay schemes, facilitate faster convergence and prevent overshooting (Ramos et al., 2024). Additionally, regularization techniques like dropout and batch normalization mitigate overfitting, while anchor box optimization enhances bounding box prediction accuracy (Garbin et al., 2020). YOLOv8 also benefits from transfer learning by employing pretrained weights, which accelerates training and improves performance on specialized datasets. These approach enriches the diversity of the training dataset, enhancing the model’s generalization ability and accelerating its learning process. On the other hand, Mask R-CNN’s approach may not benefit to the same extent from such complex augmentation strategies.

The loss functions used in YOLOv8 are generally more direct and computationally efficient (Zhan et al., 2022).These functions are designed to provide clearer gradients for optimization, making it easier to navigate the complex loss landscape. In contrast, Mask R-CNN employs multiple loss components (for bounding boxes, class labels, and masks), which complicates the optimization process and slows down convergence (He et al., 2017).

In essence, the faster convergence of YOLOv8 is not just the result of one factor, but a combination of network architecture and training strategies, which are intertwined to form an efficient and powerful system. The redesigned structure, innovative augmentation techniques, and simplified loss functions all make YOLOv8 an important method in scenarios where fast training is critical.

### 4.2 Effect of Super-Resolution Reconstruction on the YOLOv8

Super-resolution techniques typically enhance image details such as textures and edges, which are not particularly crucial for object detection tasks where YOLOv8 relies more on macro features like shape, contours, and color distribution. Additionally, the increased image resolution resulting from SR reconstruction imposes a heavier computational burden during training. The larger input size significantly increases the computation required for convolution operations, leading to extended training times and potentially affecting the model’s training efficiency. In the studies conducted by Shahi et al., 2023 and Genze et al., 2022 the challenge of low-resolution UAV images was addressed by lowering the flight altitude to 5 meters and investing in high-resolution cameras. However, drones cannot execute automatic flight plans effectively at such a low altitude, making it difficult to create orthomosaics and eliminate image distortion. Furthermore, high-resolution cameras are prohibitively expensive, limiting their practicality for widespread use.

Our study explored an alternative approach using super-resolution (SR) reconstruction technology to enhance the training set for YOLOv8. Contrary to expectations, the accuracy of YOLOv8 did not improve significantly with SR reconstruction. This may be attributed to the introduction of artifacts or noise by the super-resolution algorithm, as noted by Wang et al., (2021). These artificial details can confuse the model, hindering its ability to learn effective features and negatively impacting accuracy.

YOLOv8 is meticulously optimized for standard image resolutions. To fully leverage the benefits of super-resolution images, different network structures or parameter tuning may be necessary. Without re-optimization for SR images, the model may fail to capitalize on the enhanced features.

This study reveals the limitations of SR reconstruction in improving YOLOv8’s performance for weed segmentation in UAV images. Future work could focus on developing SR algorithms that minimize artifacts or exploring other image enhancement techniques. Additionally, fine-tuning the network architecture and parameters for SR images could further investigate the potential benefits. Conclusively, while SR reconstruction presents an innovative approach to overcoming resolution challenges, its current application in YOLOv8 for UAV-based weed segmentation requires further refinement to realize its full potential.

### 4.3 Performance of YOLOv8 in the Presence of Multiple Types of Weeds

It is common that different species weeds grow in the same field, thus it is critical to detecting multiple weed species.As expected, the segmentation model of YOLOv8 has a decreased accuracy in the presence of multiple types of weeds, which may be attributed to fact that weed species may have a high degree of visual similarity, especially in features such as shape, color, and texture (Genze et al., 2022). This similarity can make it difficult for the model to distinguish between different types of weeds, resulting in decreased accuracy in classification and segmentation.

If the types of weeds and scenes in the training data are complex and diverse, it may be difficult for the model to learn enough features to accurately distinguish each weed. Complex backgrounds and overlapping plants can also increase the difficulty of the segmentation task (Shahi et al., 2023). In the training data, the number of samples of different types of weeds may be unbalanced. Some types of weeds may appear less frequently, which will cause the model to perform poorly on these rare types because the model does not have enough opportunities to learn the features of these types during training.

The segmentation task requires classifying each pixel in the image, which is more fine-grained than simple object detection. The small parts and complex morphology of the weeds require the model to have higher resolution and fine feature extraction capabilities, which increases the difficulty of the task.

It may also be that due to insufficient data augmentation and preprocessing, the model may not be able to fully learn the diverse features of weeds in different environments and conditions.

### 4.4 Effects of Weed Control on Yield

The significant increase in potato yields due to chemical and mechanical weed control is because weeds compete with crops for water, nutrients and light resources in the soil. Through chemical treatment, the number of weeds is reduced, so that more resources can be used by crops, thereby promoting healthy crop growth and increased yields. Weeds are often hosts for some pests and pathogens (Storkey and Westbury, 2007). Through effective weed control, the habitat of these harmful organisms can be reduced, the probability of pests and diseases can be reduced, and the health of crops can be further protected (Storkey and Westbury, 2007).

In Figure 14, the yield data distribution under chemical weed control is more stable than that under mechanical weed control, which may be because chemical herbicides can be widely used on different types of weeds and provide comprehensive and consistent weed control effects (Kraehmer et al., 2014). Many chemical herbicides have a broad spectrum of weed control and can effectively control a variety of weed species, while mechanical weed control is usually more dependent on physical location and accuracy of operation, and the effect may not be as consistent as chemical methods (Paul et al., 2024). Chemical herbicides often have a certain residual effect period, which can continuously inhibit the regeneration and germination of weeds. Mechanical weed control only physically removes existing weeds and has no lasting control effect on future new weeds. Frequent operation may be required to maintain weed control effects. The application process of chemical weed control is relatively simple, mainly relying on spraying equipment. The operation process is highly standardized, reducing the uncertainty and variability of manual operation. Mechanical weed control, on the other hand, requires consideration of the operation, maintenance and on-site conditions of mechanical equipment. The operation is highly variable, and the effect is easily affected.

### 4.5 Limitations and future work

Although the weed segmentation model in this study performs well, there are still many limitations. For example, deep learning models usually require a large number of high-quality, accurately annotated datasets for training. Acquiring and annotating this data (especially pixel-level annotation) requires a lot of manpower and time, which is particularly difficult for weed segmentation tasks because weeds are numerous and have different forms. In the future, we will try to use the field image dataset to train the Real-ESRGAN model specifically for super-resolution reconstruction of field drone images, and explore whether this model can truly improve the quality of the dataset and help deep learning research in agriculture.

Weeds and crops may have highly similar visual features, such as color, shape, and texture. This makes it more difficult for the model to distinguish between the two, resulting in misclassification and segmentation errors. The farmland environment is complex and changeable, and different lighting, shadows, soil backgrounds, climatic conditions, etc. will affect the quality and characteristics of the image. After the model is trained in one environment, it may be difficult to generalize to other different environments. The training dataset may have uneven sample distribution problems, and some weed species may appear less frequently in the dataset, resulting in poor performance of the model on these weeds. The imbalanced ratio of crops to weeds in the dataset may cause the model to be more inclined to identify common species, while the recognition effect of rare species is poor. We will explore the use of more advanced network architectures, loss functions, and training strategies to improve the accuracy and generalization ability of the model, and further optimize the model for different environmental conditions and weed species.

Integrating deep learning models into existing agricultural equipment and management systems may face technical and operational challenges. In particular, it may be difficult to integrate and maintain deep learning systems in areas with limited resources and insufficient technical support. In practical applications, problems such as poor data quality, unstable network connections, and hardware device failures may also be encountered, which will affect the effectiveness and reliability of deep learning models. We will also explore the integration of deep learning models with agricultural equipment and management systems and develop more practical and efficient weed management solutions. We will consider the development of real-time processing systems and explore the use of edge computing and IoT technologies to achieve real-time weed detection and control. Solving these problems requires continuous effort in collecting high quality training data and annotation, model optimization, computing resources, environmental adaptability, real-time processing, and interpretability.

## 5 Conclusion

In summary, this study verified our hypothesis that super-resolution reconstruction can help SAM improve the quality of annotation and thus improve the model accuracy by improving the resolution of the image. The results show that Mask R-CNN and YOLOv8 have excellent weed segmentation effects, but when multiple types of weeds exist, the accuracy of the model decreases slightly. By comparison, it is found that Mask R-CNN has higher accuracy, but the YOLOv8 model converges faster. In addition, it also verified that it is feasible to apply the YOLOv8 segmentation model to yield prediction. Besides, this study explored the effects of weeds and nitrogen fertilizer on potato yield and the application of deep learning models in weed segmentation. The main findings showed that different nitrogen fertilizer treatments did not significantly affect yield, but different weed treatments did, while there was no significant difference between chemical and mechanical weeding. The effect of chemical treatment was more consistent, making the yield more stable.

Future studies should consider larger sample sizes and include more accurate annotation techniques to further validate these findings. Overall, this study deepens our understanding of weed segmentation using drone orthomosaic images and deep learning. This study provides a reference for further development of weed segmentation models and contributes to the training data pool of drone-based weed images, and thus has great practical value for precise weed control.

## 6 Acknowledgements

The author sincerely thanks the Field Crop Unit for their support in conducting the field trials.

## 7 CRediT authorship contribution statement

Chenghao Lu: Conceptualization, Data curation, Formal analysis, Investigation, Methodology, Software, Validation, Visualization, Writing – original draft.

Kang Yu: Conceptualization, Funding acquisition, Project administration, Resources, Supervision, Writing – review and editing.

## 9 Data availability statement

The data supporting the conclusions of this article will be made available on the public repository.

